# Microglia at Sites of Atrophy Restrict the Progression of Retinal Degeneration via Galectin-3 and Trem2 Interactions

**DOI:** 10.1101/2023.07.19.549403

**Authors:** Chen Yu, Eleonora M Lad, Rose Mathew, Sejiro Littleton, Yun Chen, Kai Schlepckow, Simone Degan, Lindsey Chew, Joshua Amason, Joan Kalnitsky, Catherine Bowes Rickman, Alan D Proia, Marco Colonna, Christian Haass, Daniel R Saban

## Abstract

Degenerative diseases of the outer retina, including age-related macular degeneration (AMD), are characterized by atrophy of photoreceptors and retinal pigment epithelium (RPE). In these blinding diseases, macrophages are known to accumulate ectopically at sites of atrophy, but their ontogeny and functional specialization within this atrophic niche remain poorly understood, especially in the human context. Here, we uncovered a transcriptionally unique profile of microglia, marked by galectin-3 upregulation, at atrophic sites in mouse models of retinal degeneration and in human AMD. Using disease models, we found that conditional deletion of galectin-3 in microglia led to defects in phagocytosis and consequent augmented photoreceptor death, RPE damage and vision loss, suggestive of a protective role.

Mechanistically, Trem2 signaling orchestrated the migration of microglial cells to sites of atrophy, and there, induced galectin-3 expression. Moreover, pharmacologic Trem2 agonization led to heightened protection, but only in a galectin-3-dependent manner, further signifying the functional interdependence of these two molecules. Likewise in elderly human subjects, we identified a highly conserved population of microglia at the transcriptomic, protein and spatial levels, and this population was enriched in the macular region of postmortem AMD subjects. Collectively, our findings reveal an atrophy-associated specialization of microglia that restricts the progression of retinal degeneration in mice and further suggest that these protective microglia are conserved in AMD.

**One Sentence Summary:** A common neuroprotective response of microglia at the site of retinal atrophy is identified in mice and humans.

## INTRODUCTION

Microglia, the resident macrophages of the central nervous system (CNS) (*1–5*), are highly specialized to respective microenvironments such that their functionality can vary by location and pathological perturbation, as shown in the retina and elsewhere in the CNS (*6–10*). This distinctive adaptability is pertinent in neurodegenerative states as well, where microglia migrate to the area of CNS pathology and alter their molecular and functional profiles (*6, 11–13*). There is also an intrinsic microglial contribution in producing these changes (*14–16*), since infiltrated monocyte-derived macrophages do not fully adopt the same characteristics (*17–21*). Hence, both ontogeny and location are critical factors in understanding microglial roles in neurodegenerative diseases.

Degenerative diseases of the outer retina, including age-related macular degeneration (AMD), are common causes of blindness in adults and are characterized with retinal atrophy of photoreceptors and retinal pigment epithelium (RPE). AMD alone afflicts approximately 196 million worldwide (*22*), yet only 10% of these cases are treatable, highlighting a major unmet medical need (*23*). Innate immunity is deemed important in AMD pathobiology (*24–27*), and the major genetic risk genes for AMD (*28*), including *CFH*, *ARMS2-HTRA1*, *APOE*, and *C3*, are all expressed by or impact the immune system (*29–32*). Significant for these diseases, macrophages migrate and accumulate ectopically in the subretinal space, the area adjacent to the sites of atrophy (*33–37*). However, whether the cells are microglia-derived is incompletely known, particularly in the human context. Moreover, while neuroinflammation is generally thought to contribute to the disease process, the profile and function of these ectopic subretinal immune cells are largely unknown. Here, we set out to elucidate the core phenotypic and functional signature of microglia in degeneration with a focus on the subretinal space, the mechanisms underlying their contributions to disease, and the potential of targeting these cells pharmacologically.

## RESULTS

### Identification of a common transcriptional signature of subretinal microglia

To investigate the responses by microglia in outer retinal degeneration, we performed single-cell RNA-sequencing (scRNA-seq) of CD45^+^ cells purified from mouse retinas in four distinct model settings. These included i) 2-month-old wild type (WT) mice (naïve) as a young adult baseline; ii) *Rho^P23H/+^* (P23H) knock-in mice as a genetic model of photoreceptor degeneration (*38*); iii) sodium iodate (NaIO_3_) model of acute injury to the RPE (*39*); iv) and 2-year-old WT mice to mimic advanced aging (**Fig. S1A** and **S1B**). Analysis of these settings identified eight clusters of major immune cell populations, including microglia, monocyte-derived macrophages, perivascular macrophages, monocytes, T cells, B cells, natural killer cells, and neutrophils (**Fig. S1C**).

Next, to profile these retinal microglial clusters, we integrated our published dataset of Cx3cr1^+^ sorted retinal cells from mice subjected to light damage (LD) (*6*), an acute model of photo-oxidative stress induced photoreceptor degeneration. Successful integration of this dataset was established by the presence of overlapping clusters of microglia, perivascular macrophages, monocyte-derived macrophages, as well as contaminant retinal neurons, but not other immune cells (**Fig. 1A**). Integrated analysis of over 15,000 macrophages including microglia revealed comparable cluster types among the four degeneration settings (**Fig. 1B**). The cluster of subretinal microglia, as previously identified in the LD model (*6*), was found in samples primarily from all four degeneration models but not from naïve retinas (**Fig. 1C**). Differential gene analysis revealed a core transcriptional signature for subretinal microglia that was common among four degeneration models (**Fig. 1D**). The top shared upregulated genes included *Lgals3, Cd68, Gpnmb, Fabp5, Vim*, *Cstb* and *Cd63*, while downregulated genes included homeostatic microglial markers, such as *P2ry12, Tmem119,* and *Cx3cr1* (**Table S1**).

**Fig. 1.**
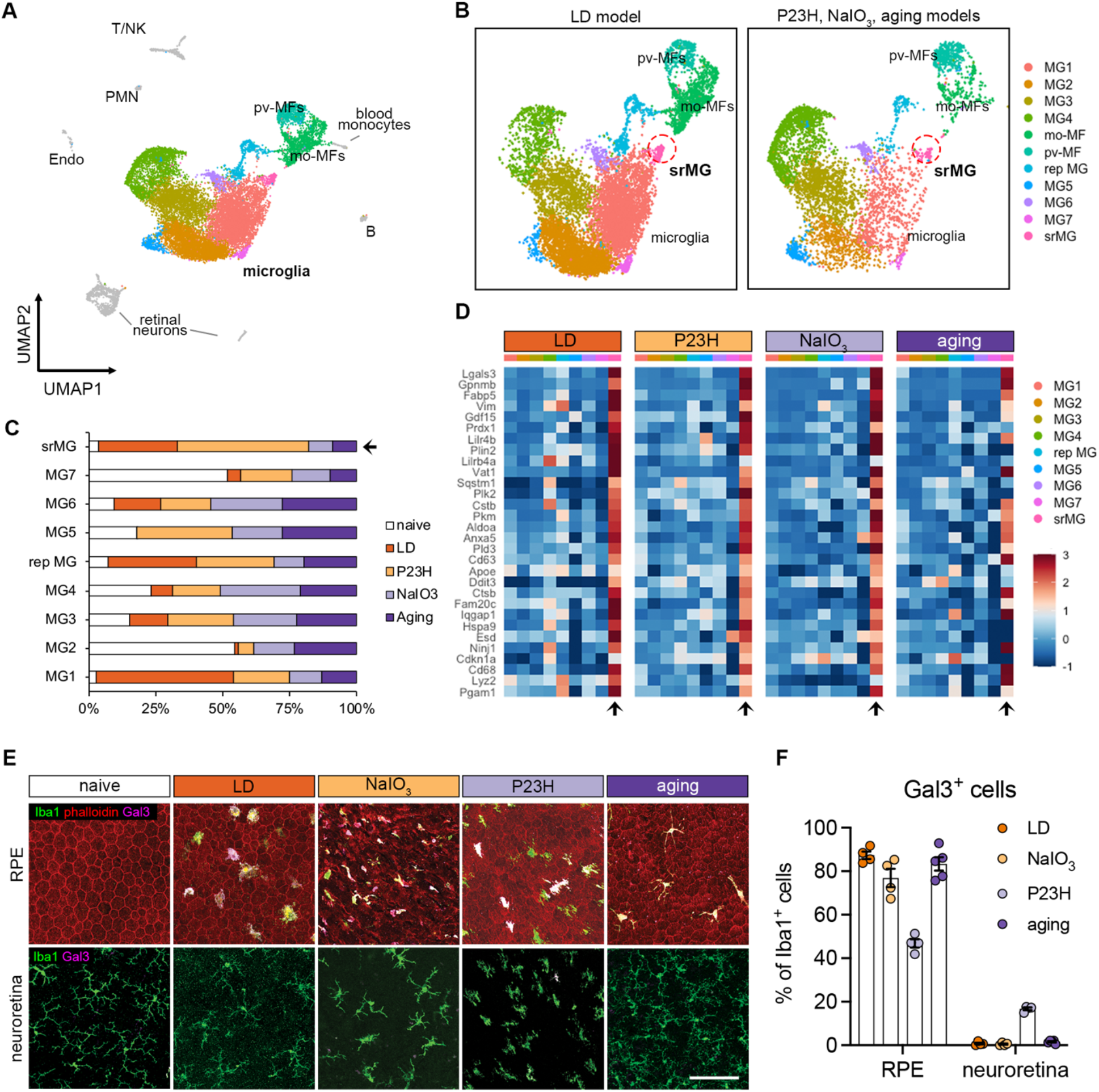
The microglial population present in the subretinal space shares a common signature in mouse models of photoreceptor degeneration, RPE degeneration, and advanced aging. (**A**) UMAP plot showing integrated clustering of immune cells samples from four mouse models of retinal degeneration, including LD model (sorted by Cx3cr1^+^), NaIO_3_ model (CD45^+^), P23H model (CD45^+^) and aging model (CD45^+^) and naïve mice (CD45^+^). A total of 15,623 macrophages, including 13,489 microglia, were integrated among four models. PMN, polymorphonuclear neutrophils; mo-MFs, monocyte-derived macrophages; pv-MFs: perivascular macrophages; NK, natural killer. (**B**) UMAP plots showing integrated macrophage clusters by two datasets. Dash circles indicate subretinal microglia (srMG). (**C**) Percentage of sample distribution by clusters. The arrow indicates the enrichment of srMG cluster from degenerating retinas. (**D**) Heatmap of top 30 conserved marker genes of subretinal microglia shared by each model across clusters. Genes were ranked by fold changes. Arrows indicate srMG cluster. (**E**) *In situ* validation of Gal3 expression on the apical RPE (top) or in the neuroretina from the inner plexiform layer (bottom). Iba1 (green), phalloidin (red, only in RPE) and Gal3 (magenta). Scale bar: 100μm. (**F**) Percentage of Gal3^+^ cells relative to Iba1^+^ cells between RPE and neuroretina tissues across models.

As our integrated scRNA-seq analysis revealed that *Lgals3* expression was highly enriched in the subretinal microglia cluster only (**Fig. 1SD)**, we further analyzed its expression in the disease models. We examined the co-expression of galectin-3 (Gal3) protein *in situ* and Iba1 in all four models and compared it to naïve mice (**Fig. 1E**). We noted that the morphology of subretinal Iba1^+^ is different among models (**Fig. S1E** and **S1F**), which is likely due to the distinct features and disease progressions in the individual models. However, we observed Gal3^+^ Iba1^+^ cells in all four models predominantly located in the subretinal space, on the apical aspect of the RPE. Few to none of Gal3^+^ Iba1^+^ cells were detected in the plexiform layers of neuroretina or the RPE of naïve retina (**Fig. 1E** and **1F**), suggesting that induction of *Lgals3* upregulation occurs in the subretinal space. Together, our findings suggest the presence of Gal3^+^ subretinal microglia with a common transcriptional signature in distinct forms of retinal degeneration and advanced aged mice.

### Deletion of galectin-3 in subretinal microglia exacerbates retinal degeneration

Previously, we showed that depletion of endogenous microglia led to excessive accumulation of photoreceptor debris and massive structural damage to the RPE (*6*). Here, we investigated whether Gal3 mediates this disease-restricting microglial response in retinal degeneration. We began by analyzing global *Lgals3* knockout (KO) mice and confirmed that young adult naïve *Lgals3* KO mice at the age of 2 months had normal RPE morphology and marginal subretinal Iba1^+^ cells (**Fig. S2A**), consistent with a previous report (*40*). Next, we subjected young *Lgals3* KO and age-matched WT mice to the LD model and compared their retinal phenotypes. *Lgals3* KO mice had substantially increased dysmorphic RPE and elevated TUNEL^+^ cells in photoreceptor layers compared with WT (**Fig. 2A-2D**), phenocopying the microglia depletion setting in LD shown in a prior study (*6*). To assess if Gal3 has an impact on phagocytosis by subretinal microglia, we examined the rhodopsin level in the subretinal Iba1^+^ cells. The results showed that *Lgals3* KO mice failed to engulf dead photoreceptors (**Fig. 2E**) and showed a dramatic drop of rhodopsin^+^ subretinal microglia and a massive accumulation of photoreceptor debris (**Fig. 2F**), implicating a critical role of Gal3 in the clearance of dead photoreceptors. The augmented damage could not have been due to subretinal Iba1^+^ cell quantity, as their densities were comparable in both groups (**Fig. S2A** and **S2B**). Next, we compared these mice in the advanced aging setting at the age of 2 years. Aged *Lgals3* KO mice showed increased RPE size and reduced visual function measured by electroretinograms (ERG) with attenuated scotopic a-waves and b-waves (**Fig. 2G-2I**). Finally, we bred *Lgals3* KO onto P23H mice, a clinically relevant mouse model of retinitis pigmentosa (*38*). Despite comparable subretinal Iba1^+^ cell numbers (**Fig. S2C** and **S2D**), loss of Gal3 led to electrophysiological deficits (**Fig. 2J, Fig. S2E**) and augmented thinning of photoreceptor layers (**Fig. 2K, Fig. S2F**) in P23H mice. Similar phenotypes of *Lgals3* KO mice were also recently reported in NaIO_3_ model (*41*). Collectively, these data demonstrate that Gal3 plays a protective role in different forms of retinal degeneration.

**Fig. 2.**
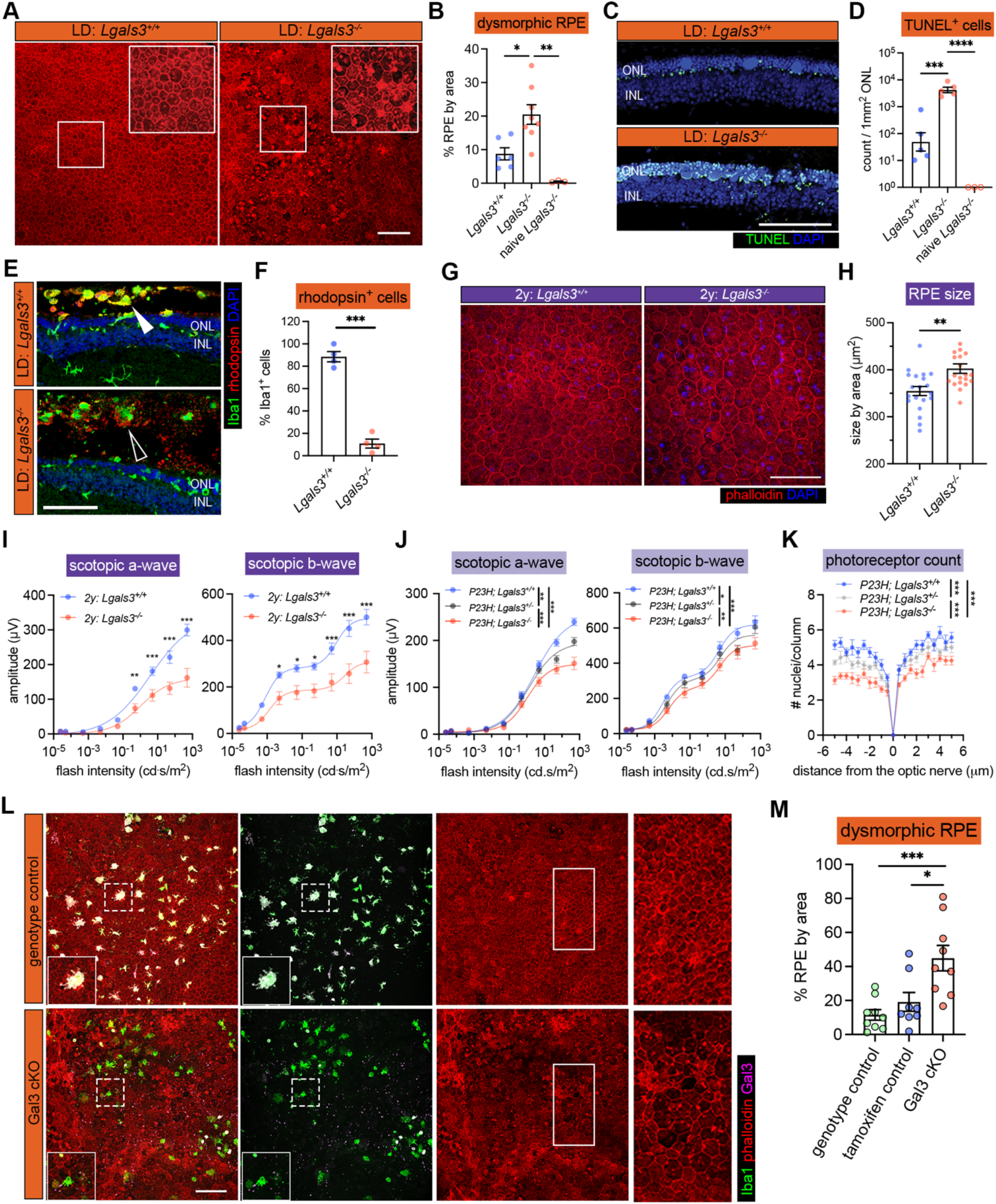
Galectin-3 expressed by subretinal microglia is central in restricting disease progression in acute, genetic, and aging mouse models of retinal degeneration. (**A**) Images of phalloidin staining in WT and *Lgal3^-/-^* RPE tissues in LD. (**B**) Quantifications of dysmorphic RPE cells (n=6, 7 and 3, respectively). (**C**) TUNEL (green) and DAPI (blue) staining in WT and *Lgal3^-/-^* retinal cross sections in LD. ONL and INL, outer and inner nuclear layers. (**D**) Quantifications of TUNEL^+^ photoreceptors in ONL (n=5, 5 and 3, respectively). (**E**) Rhodopsin (red) and Iba1 (green) staining in WT and *Lgal3^-/-^* retinal cross sections in LD. Images from single planes of confocal scans were shown. (**F**) Quantifications of rhodopsin+ subretinal microglia (n=4 per group). (**G**) Images of phalloidin staining in WT and *Lgal3^-/-^* RPE tissues at 2 years of age. (**H**) Quantifications of RPE cell size. Dots represent individual images with n=5 mice per group. (**I**) ERG data showing scotopic a- and b-waves in 2-year-old WT (n=5) and *Lgal3^-/-^* (n=5) mice. (**J**) Scotopic a- and b-waves of ERG data among *Lgal3^+/+^* (n=12), *Lgal3^+/-^*(n=6) and *Lgal3^-/-^* (n=10) in P23H mice. (**K**) Quantifications of ONL thickness among *Lgal3^+/+^* (n=7), *Lgal3^+/-^* (n=7), and *Lgal3^-/-^* (n=8) in P23H mice. (**L**) Representative images of dysmorphic RPE cells in Gal3 cKO in LD. Iba1, green; phalloidin, red; Gal3, magenta. (**M**) Quantifications of dysmorphic RPE cells in Gal3 cKO mice (n=9) compared with genotype control (*Cx3cr1^CreER/+^Lgals3^fl/fl^* mice, n=9) and tamoxifen control (*Cx3cr1^CreER/+^* mice treated with tamoxifen, n=8). Scale bars: 100μm. Data were collected from 2-3 independent experiments. *: p<0.05; **: p<0.01; ***: p<0.001. One-way ANOVA with Tukey’s post hoc test (B, D and M); unpaired Student’s t-test (F and H); two-way ANOVA with Tukey’s post hoc test (I, J and K).

We next wanted to determine the microglia specific role of Gal3 *in vivo*, as this protein can also be expressed by monocyte-derived cells, Müller cells and astrocytes. We bred *Lgals3^fl/fl^* mice (*42*) onto a C57BL/6J background and then crossed these mice with *Cx3cr1^CreER^* for a microglial conditional KO (cKO). We included the genotype control (*Cx3cr1^CreER/+^; Lgals3^fl/fl^* without tamoxifen) and tamoxifen control (*Cx3cr1^CreER/+^* with tamoxifen) groups. Considering the critical role of Cx3cr1 in regulating microglial function and sensitivity to light-induced retinal degeneration (*33, 43*), we only used *Cx3cr1^YFP-CreER^* heterozygous mice hereafter and tested subjecting these mice to the LD model. We achieved ∼75% deletion efficiency of Gal3 in subretinal microglia (**Fig. 2L, Fig. S2G**). Gal3 cKO led to increased dysmorphic RPE cells and no change in the densities of subretinal Iba1^+^ cell compared with control mice (**Fig. 2M, Fig. S2H**), consistent with the findings of global *Lgals3* KO mice. Collectively, these results demonstrate that Gal3 contributes to the protection by subretinal microglia but is not required for microglial migration.

### Trem2 regulates microglial subretinal migration and galectin-3 expression

Our integrated scRNA-seq analysis revealed the upregulation of genes associated with trigger receptor expressed by myeloid cell 2 (Trem2) signaling in subretinal microglia, including *Syk*, *Apoe,* and *Ctnnb1* (**Fig. 3A**). We also found that protein levels of Trem2 and its downstream effector tyrosine kinase Syk were dramatically increased in subretinal microglia (**Fig. 3B** and **3C**), although Trem2 mRNA level remained relatively unchanged in scRNA-seq (**Fig. 3A**). Based on a recent study showing Gal3 as a novel ligand of Trem2 (*44*), we tested the hypothesis that Trem2 regulates Gal3-mediated function by subretinal microglia. Following this lead, we co-immunolabelled Iba1, Trem2, and Gal3 on retinas from LD-subjected mice, where we previously demonstrated the predominance of microglia in subretinal space (*6*). Our results revealed the colocalization of Trem2 and Gal3 in subretinal microglia (**Fig. 3D**). Specifically, Trem2 and Gal3 were colocalized on the surface side of subretinal microglia facing the apical aspect of RPE (**Fig. 3D**, **Fig. S3A**). Hence, our results implicate a functional specification of Trem2-Gal3 interactions by subretinal microglia in restricting RPE injury and other aspects of retinal degeneration.

**Fig. 3.**
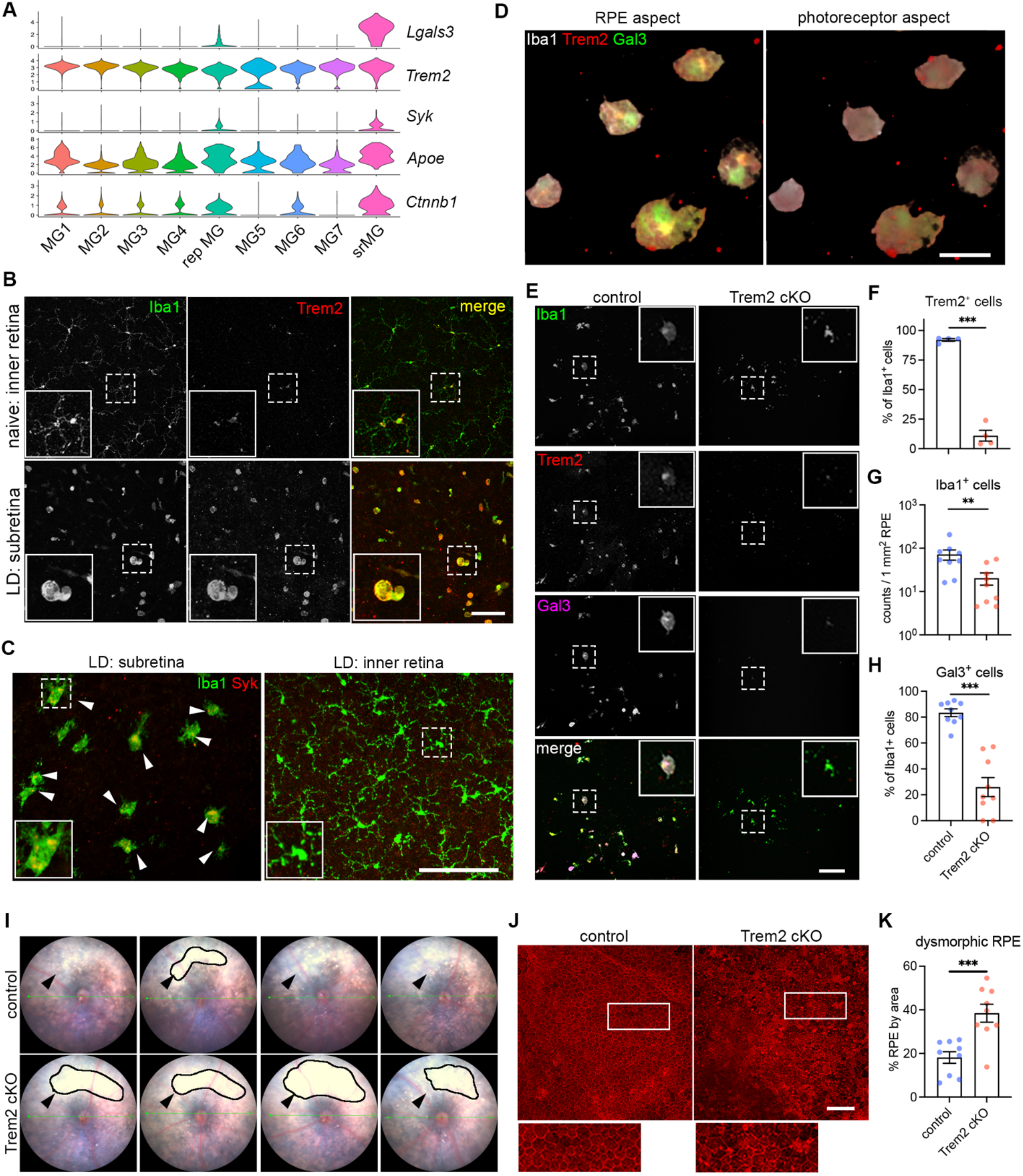
Trem2 regulates microglial migration and promotes galectin-3-mediated protection. (**A**) Violin plots showing the upregulation of genes (*Lgals3, Syk* and *Ctnnb1I)* related to Trem2 signaling by subretinal microglia from the integrated dataset of all four mouse models. (**B**) Images of Iba1 (green) and Trem2 (red) staining in naïve microglia from inner retina and subretinal microglia in LD. (**C**) Images of Iba1 (green) and Syk (red) staining in subretinal microglia and microglia from inner retina in LD. (**D**) 3D rendering images of Gal3 (green), Trem2 (red) and Iba1 (white) staining in subretinal microglia in LD. Views from both the apical RPE aspect and neuroretina aspect are shown. (**E**) Images of Iba1 (green), Trem2 (red) and Gal3 (magenta) staining in subretinal microglia between control and Trem2 cKO mice in LD. (**F-H**) Quantifications of Trem2 depletion (F, n=4 per group), Iba1^+^ cells (G, n=9) and Gal3^+^ cells (H, n=9) between control and Trem2 cKO mice. (**I**) Fundus images showing increased subretinal white lesions in of Trem2 cKO mice in LD as indicated by arrows. Images from four individual mice per group are shown. (**J**) Images of phalloidin staining in RPE tissues from control and Trem2 cKO mice in LD. (**K**) Quantifications of dysmorphic RPE cells between control and Trem2 cKO mice (n=9 per group). Scale bars: 50μm (D); 100μm (B, C E,and J). Data were collected from 2 independent experiments. **: p<0.01; ***: p<0.001. Unpaired Student’s t-test (F-H).

To examine whether Trem2 mediates the protective microglia response, we pharmacologically inhibited Trem2 signaling using anti-Trem2 mAb178 via tail vein injection in mice before LD exposure. This antibody blocks the ligand binding of lipidated HDL and Trem2 signaling (*45, 46*). Our results showed that Trem2 blockade led to increased subretinal white lesions (**Fig. S3B**). However, unlike *Lgals3* deletion, Trem2 blockade led to a reduction of subretinal Iba1^+^ cells (**Fig. S3C** and **S3D**), suggesting that Trem2 mediates migration of microglia to the subretinal space (*47–49*). The observed reduction was accompanied with reduced Gal3 expressing Iba1^+^ cells in the subretinal space and amplified RPE dysmorphogenesis (**Fig. S3E-S3G**), therefore linking Gal3 and Trem2 activity with microglial-mediated neuroprotection.

We next examined if this Trem2-mediated response is microglia-specific by crossing *Cx3cr1^CreER^* with *Trem2^fl/fl^* mice to achieve microglial *Trem2* cKO. With an 85% Trem2 deletion efficiency in retinal microglia (**Fig. 3E** and **3F**, **Fig. S3H**), LD-subjected Trem2 cKO mice had reduced subretinal Iba1^+^ cells and Gal3 expression (**Fig. 3G** and **3H**). Subretinal microglia in these *Trem2* cKO mice also appeared morphologically less distended (**Fig. 3E**), which is consistent with reduced phagocytic activity (*49–51*). Corroborating results with pharmacologic Trem2 blockade, we observed enhanced subretinal white lesions via fundoscopy (**Fig. 3I**) and increased RPE damage (**Fig. 3J** and **3K**) in *Trem2* cKO relative to controls in the LD model. We therefore conclude that Trem2 is critical in microglial-mediated protection by regulating subretinal migration and inducing Gal3 expression.

### Augmentation of Trem2 activity ameliorates retinal degeneration in a Gal3-dependent manner

Elevated levels of soluble TREM2 were observed in cerebrospinal fluid of Alzheimer’s patients (*52, 53*). To determine whether increased levels of soluble Trem2 are also found in response to retinal degeneration, we measured soluble Trem2 in vitreous and retinal fluids collected from LD-subjected mice versus naïve controls. We found substantially increased soluble Trem2 levels in LD fluid samples compared with those collected from naïve controls (**Fig. 4A**). In contrast, *Trem2* cKO mice subjected to LD had a reduced level of soluble Trem2, indicating a microglia-derived origin (**Fig. 4A**). Therefore, these findings support our observations on increased Trem2 expression and previous reports of elevated α-secretases, including ADAM10/17 in retinal degeneration as well (*54, 55*).

**Fig. 4.**
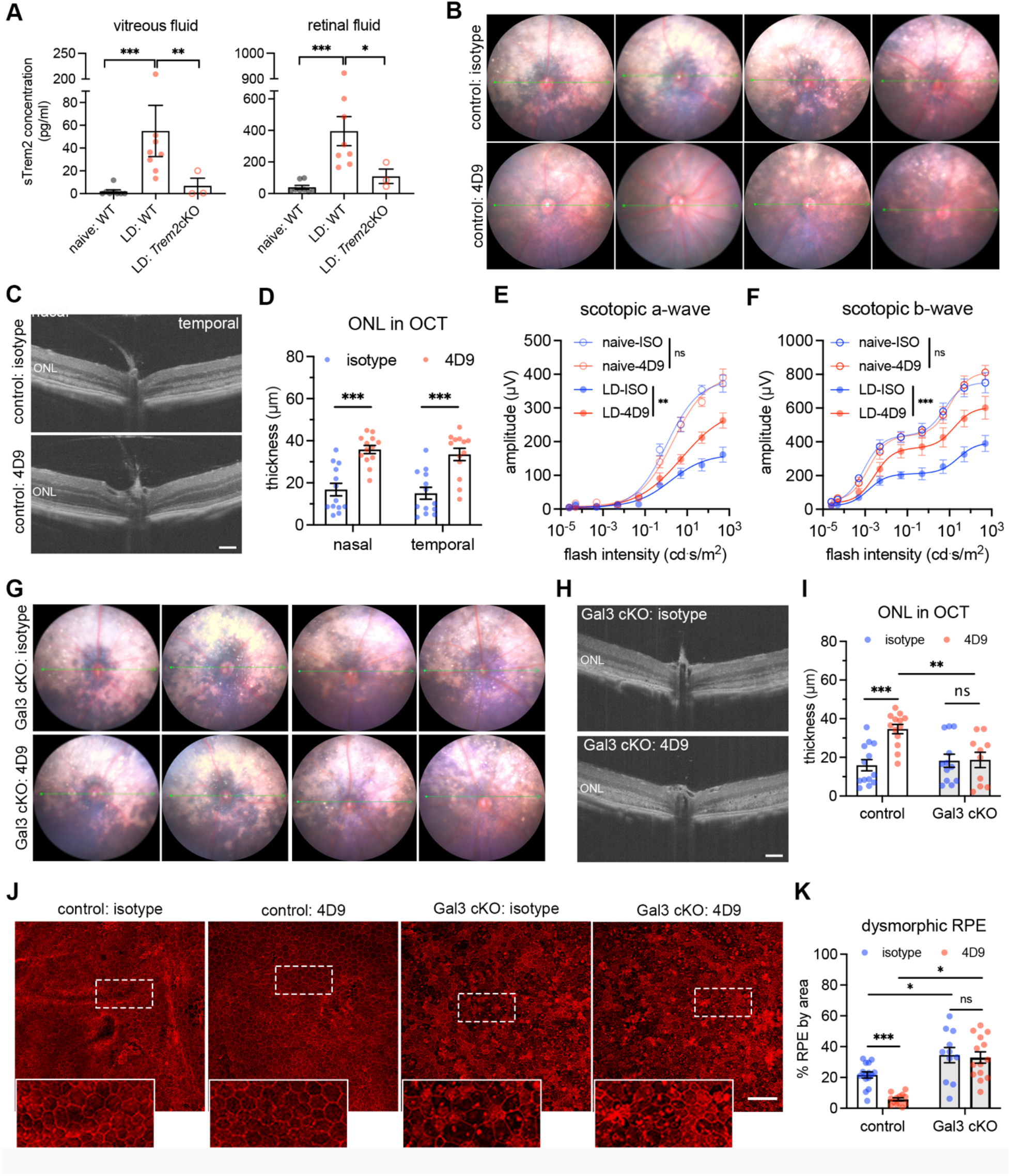
Bolstering galectin-3-dependent Trem2 signaling by microglia prevents retinal degeneration. (**A**) ELISA of soluble Trem2 (sTrem2) in vitreous fluid and retinal fluid from naïve WT mice, WT and Trem2 cKO mice subjected to LD. (**B**) Fundus images of mice treated with isotype control or 4D9 anti-Trem2 in LD. Four individual mice per group are shown. (**C**) Representative OCT images of mice treated with isotype or 4D9 in LD. (**D**) Quantifications of outer nuclear layer (ONL) thickness by OCT (n=13 per group). ONL thickness was measured at both nasal and temporal sides. (**E** and **F**) Scotopic a-waves and b-waves of ERG data among mice treated with isotype or 4D9 in naïve or LD setting (n=5 per group). (**G**) Fundus images of Gal3 cKO mice treated with isotype or 4D9 in LD. Four individual mice per group are shown. (**H**) Representative OCT images of Gal3 cKO mice treated with isotype control or 4D9 anti-Trem2 in LD. (**I**) Quantifications of average ONL thickness by OCT between control and Gal3 cKO mice treated with either isotype or 4D9 (n=13 per group). (**J**) Images of phalloidin staining of control and Gal3 cKO RPE treated with isotype or 4D9 in LD. (**K**) Quantifications of dysmorphic RPE cells (n=15, 13, 11 and 13, respectively). Scale bars: 100μm. Data were collected from 2-4 independent experiments. *: p<0.05; **: p<0.01; ***: p<0.001. Unpaired Student’s t-test (F-H). One-way ANOVA with Tukey’s post hoc test (A); two-way ANOVA with Tukey’s post hoc test (D-F, I and K).

To test the hypothesis that a Trem2 gain-of-function would further promote its activity and thereby lead to additive protection, we leveraged the dual function 4D9 anti-Trem2 antibody. This antibody binds the Trem2 stalk region recognized by ADAM10/17 to impede ectodomain shedding, cross-link and stabilize Trem2 expression on microglial surface, thereby promoting phospho-Syk signaling (*56*). Specifically, we examined whether 4D9 can ameliorate degeneration outcomes in the LD model. We administrated one dose of 4D9 or isotype antibody to mice via tail vein injection before LD exposure. Fundoscopy showed that retinal injury was substantially reduced in 4D9 treated mice compared with isotype controls (**Fig. 4B**). Likewise, optical coherence tomography (OCT) images revealed preservation of the photoreceptor layer with over 2-fold increase on average in 4D9 treated mice (**Fig. 4C** and **4D**). Consistent with the OCT changes, scotopic ERGs showed that 4D9 treatment protected visual function compared with isotype controls in degeneration (**Fig. 4E** and **4F**). In addition, the human IgG domain of 4D9 antibodies was predominantly detected in subretinal microglia but was rarely observed in Iba1^+^ cells within the inner retina (**Fig. S4A-S4C**), implicating the added protection was derived from subretinal microglia. Therefore, we show that pharmacologic augmentation of Trem2 activity in subretinal microglia protects the retina from degeneration.

Lastly, we sought to address whether Trem2 and Gal3 are functionally interdependent in microglial-mediated neuroprotection, by testing whether 4D9 ameliorates retinal degeneration in *Lgals3* cKO mice. Analyses using fundoscopy and OCT imaging showed the protection rendered via 4D9 treatment was lost in *Lgals3* cKO mice, as white subretinal lesions and photoreceptor thickness were indistinguishable between 4D9 and isotype (**Fig. 4G-4I**, **Fig. S4D**). We also evaluated the impact of 4D9 treatment on the preservation of RPE integrity. Our results showed that 4D9 treatment better preserved RPE morphology relative to isotype in control mice, whereas this protective effect was lost in *Lgals3* cKO mice (**Fig. 4J** and **4K**). However, the frequencies of subretinal microglia did not significantly change between isotype and 4D9 treatments (**Fig. S4E**). Together, our findings show that Trem2 protection is dependent on Gal3 expressed by subretinal microglia, thereby further supporting their functional interdependence.

### Microglia at the sites of atrophy show a conserved molecular phenotype and are enriched in the macula of AMD patients

To address the significance of our findings in the human context, we performed another scRNA-seq of myeloid cells from postmortem neuroretina and RPE/choroid tissues of eight human donors at the age of over 70, including three AMD cases (**Table S2**). Of each donor, we used one eye to evaluate retinal and choroidal pathology (*34*), and for the contralateral eye we sorted CD45^+^CD11B^+^ cells. We captured 14,873 myeloid cells that passed QC, making it a valuable resource to uncover microglial states in human retinal tissues. Similar to mouse models studied in our aforementioned experiments, our unsupervised clustering results of neuroretina and RPE/choroid tissues revealed five clusters of three major macrophage cell types, including microglia, perivascular and monocyte-derived macrophages (**Fig. 5A, Fig. S5A, Table S3**). Among these clusters, human retinal microglia specifically expressed *TMEM119, TREM2* and *CX3CR1*, while perivascular and monocyte-derived macrophages can be distinguished by the expression of *LYVE1* and *CCR2*, respectively (**Fig. 5B**).

**Fig. 5.**
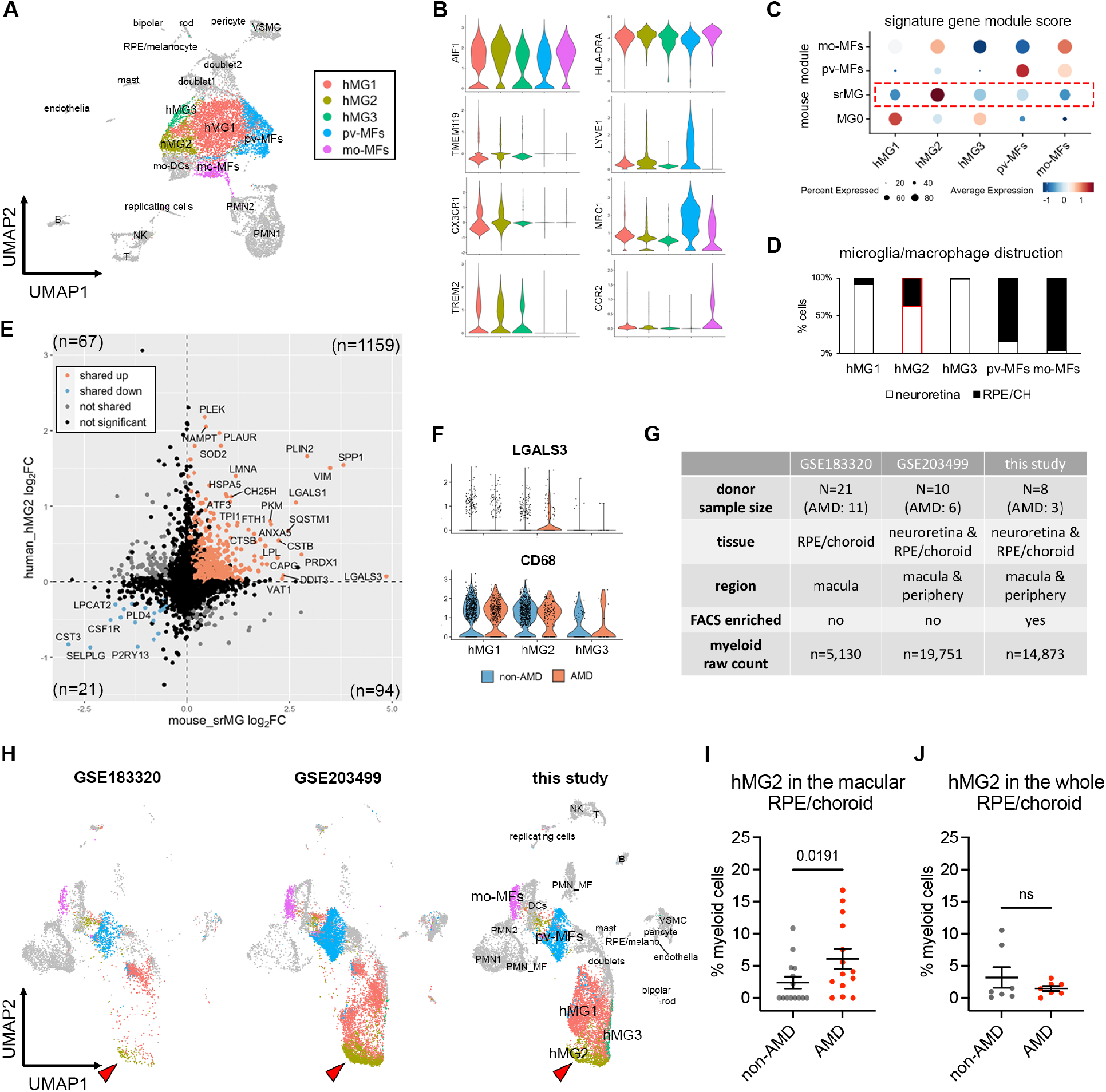
Microglia at the sites of atrophy show a conserved phenotype between mice and humans and are enriched in the macula of AMD patients. (**A**) UMAP plot showing unsupervised clustering analysis of myeloid cells from human donors. CD45^+^CD11b^+^ cells were FACS-sorted from neuroretina and RPE/choroid tissues, respectively. hMG, human microglia; mo-MFs, monocyte-derived macrophages; pv-MFs: perivascular macrophages; mo-DCs, monocyte-derived dendritic cells; VSMC, vascular smooth muscle cells. (**B**) Violin plots showing the marker expression by macrophage clusters. (**C**) Dot plots showing gene module scores of human microglia/macrophage clusters. The gene modules were generated and normalized using top 200 mouse markers from homeostatic microglia (MG0), subretinal microglia (srMG), pv-MFs and mo-MFs. (**D**) Bar graphs showing the composition of macrophage/microglia clusters by tissues. Red box indicates the enrichment of cells from RPE/choroids in hMG2 cluster. (**E**) Comparison of gene expression between mouse subretinal microglia (x axis) and human hMG2 (y axis). The number in each quadrant shows the quantity of differentially expressed genes as indicated by colors. (**F**) Violin plots showing the expression of *LGALS3* and *CD68* by microglia clusters between non-AMD and AMD donors. (**G**) Summary of three independent human AMD scRNA-seq datasets. (**H**) UMAP plots showing the label transfer of myeloid cells among datasets. Arrows indicate hMG2 clusters in each dataset. (**I** and **J**) Quantifications of hMG2 frequencies in the whole and macular RPE/choroid tissues between non-AMD and AMD donors. Mann-Whitney test (one-tailed) was used, and p-values are shown; ns: not significant.

To determine which human macrophage cluster represents subretinal microglia in AMD, we utilized the identified mouse marker genes of subretinal microglia, homeostatic microglia, and other macrophages from mice (*6*), and calculated the corresponding human gene module scores of these macrophage clusters (*57*). Two of the human macrophage clusters gained highest scores of perivascular and monocyte-derived macrophages, respectively, corroborating the identities determined above (**Fig. 5C**). By contrast, hMG2 cluster showed the highest similarity to subretinal microglia (**Fig. 5C**). Interestingly, hMG2 cluster is also composed of more cells from RPE/choroid tissues compared the other two microglia clusters (**Fig. 5D, Fig. S5B**), which is consistent with the knowledge that subretinal microglia adhere to the apical RPE, and thereby leading us to hypothesize that hMG2 cluster represents subretinal microglia in human AMD. To examine the similarities between mouse subretinal microglia and hMG2 cluster, we directly compared their gene expression profiles. Among 1,341 differentially expressed genes (DEGs) shared by mice and humans, 87.99% of these genes are similarly changed, including 1,159 upregulated genes and 21 downregulated genes (**Fig. 5E**). The pathway enrichment analysis of top shared upregulated genes inferred functions in phagocytosis, responses to oxidative stress, and lipid metabolism (**Fig. S5C**), which is line with our mouse findings. Also, *LGALS3* is enriched in cells of hMG2 clusters from AMD patients and *CD68* is expressed by all microglia clusters (**Fig. 5F**). Hence, transcriptomic analysis suggests a conserved profile for subretinal microglia between mice and humans.

To investigate whether the hMG2 cluster is associated with human AMD, we extracted myeloid cells by *AIF1* expression from another two publicly available datasets of independent AMD studies and integrated them with our dataset. The integrated dataset contains 39,754 myeloid cells collected from a total of 39 human donors with 20 AMD patients and 19 age-matched control (**Fig. 5G**). Our integrated analysis revealed similar clusters among three datasets (**Fig. S5D-S5F**), with the exception of *RHO^high^* microglia cluster exclusively derived from one AMD donor with neovascularization (**Fig. S5G1**). The GSE183320 dataset that does not contain cells from neuroretina, showed substantial decrease of microglial cells compared with the other two datasets, especially for hMG1 and hMG3, supporting a proper data integration (**Fig. 5G** and **Fig. S5F**).

To determine which cells belong to hMG2 cluster in the integrated datasets, we used two independent approaches. One approach was via labelling transfer by Seurat package (**Fig. 5H**); the second was via subclustering to determine LGALS3 expressing clusters (**Fig. S5H** and **S5I**). Both approaches showed similar results for the hMG2 frequencies. Specifically, we found that frequencies of hMG2 significantly increase in the macula of RPE/choroid tissues from AMD donors (**Fig. 5I** and **Fig. S5J**). The hMG2 cluster was also present in the elderly non-AMD group but with a lower frequency (**Fig. 5J** and **Fig. S5K**), which may represent a part of normal aging as seen in our mouse aging model. Taken together, we conclude that these subretinal microglia are most enriched in the locations associated with tissue atrophy of human AMD.

### *In situ* evidence and correlation of subretinal microglia expressing GAL3 and TREM2 in AMD subjects

Lastly, we validated the presence of these subretinal microglia in AMD by immunohistochemically staining human post-mortem tissues from another cohort of n=18 aged non-AMD and AMD donors (**Table S2**), using antibodies against human GAL3 and CD68, two markers of the subretinal microglia population that were previously validated in mice (*6*). Immunolabeling was performed on tissues sectioned through the macular region and classified according to the Sarks AMD grading scale of disease severity (*34*). Multispectral imaging revealed GAL3^+^CD68^+^ cells, stained orange-red due to co-expression, were predominantly observed in the subretinal space but rarely in the inner retina (**Fig. 6A**). The GAL3^+^CD68^+^ cells were primarily located between the neurosensory retina and the RPE, adherent to basal deposit in areas of absent RPE, and within the basal deposit between RPE and Bruch’s membrane (**Fig. SA6**). We found that GAL3^+^ CD68^+^ cells were enriched in the subretinal space within the macula in Sarks stages IV to VI (intermediate to advanced AMD), but not in aged controls represented by Sarks stages I (normal) and II (aging) or III (early AMD) (**Fig. S6A**). The double-positive cells were enriched in the regions of geographic atrophy with RPE loss (**Fig. 6B, Fig. S6B**), but also present in the areas of photoreceptor loss with preserved RPE cells, including the transitional area of AMD macula (**Fig. S6C**). Moreover, we identified a strong positive correlation between Sarks AMD grades and the frequencies of these double positive cells in the subretinal space of the macular region (**Fig. 6C**). These findings corroborate our observed enrichment of the hMG2 cluster in the macular region of human AMD subjects. In addition, we observed that our sections from both age-matched controls and AMD subjects exhibited age-related peripheral retinal degeneration, including peripheral cystoid and ischemic (paving stone) degeneration (*58–60*), wherein subretinal GAL3^+^CD68^+^ cells were also observed (**Fig. S6D**). This may partially explain the presence of hMG2 with low frequencies in aged non-AMD subjects. Of note, the frequencies of subretinal cells may be underestimated because we analyzed retinal cross-sections within the macula but not the whole macula. Taken together, these data suggest that the presence of subretinal GAL3^+^ CD68^+^ cells is a response to degeneration of the outer retina and these cells are enriched in AMD geographic atrophy.

**Fig. 6.**
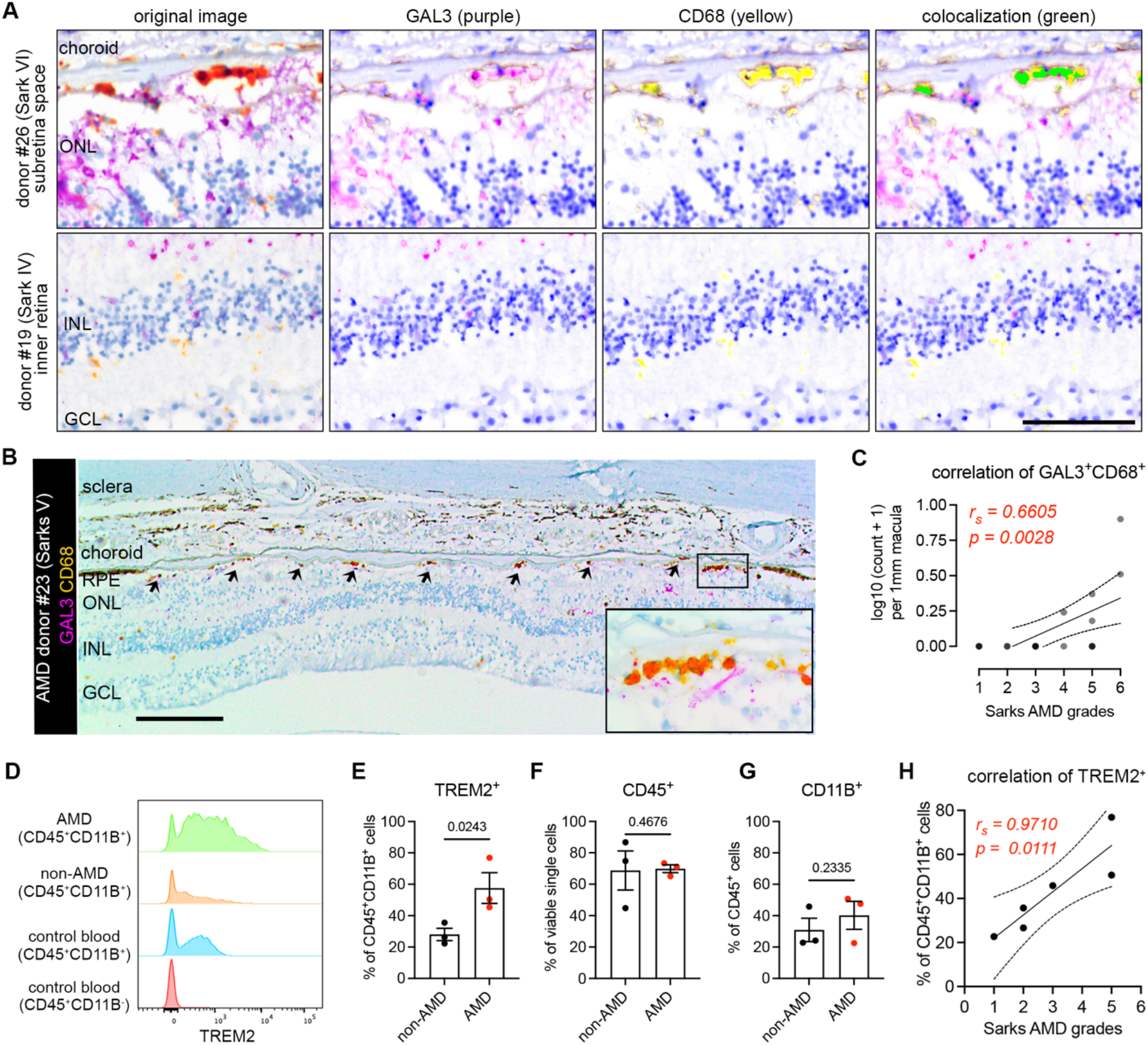
Microglia expressing galectin-3 and TREM2 are associated with AMD progression. (**A**) Multispectral imaging of GAL3 and CD68 co-staining in the subretinal space (top) and inner retina (bottom) from human donors. Unmixed purple spectrum (GAL3) and yellow spectrum (CD68) are shown. The areas of colocalized spectra are highlighted in green. Scale bar: 50μm. ONL and INL, outer and inner nuclear layers. (**B**) Representative image of Gal3 and CD68 co-staining in the macular GA region of a retinal section from an 88-year-old female donor eye with advanced AMD (Sarks V). Black insert box shows the magnification of GA with double positive cells. Scale bar: 200μm. ONL and INL, outer and inner nuclear layers; GCL, ganglion cell layer. (**C**) Correlation between the frequencies of macular Gal3^+^CD68^+^ double positive cells (y axis) and Sarks AMD grading (x axis) by Spearman’s correlation (n = 18 donors, Table S2). Coefficient and p-value are shown. (**D**) Histograms showing increased TREM2^+^ myeloid cells (CD45^+^CD11B^+^) in RPE/choroid tissues of AMD donors. Concatenated histograms were shown (n=3 per groups). Control human blood samples were used to set up flow gating. (**E-G**) Quantifications of TREM2^+^ (E), CD45^+^ (F), and CD11B^+^ (G) cell frequencies in RPE/choroid tissues between non-AMD and AMD donors. Unpaired Student’s t test is used. P-values are shown. (**H**) Correlation between the frequencies of TREM2^+^ myeloid cells (y axis) and Sarks AMD grading (x axis) in RPE/choroid tissues by Spearman’s correlation. Coefficient and p-value are shown.

To determine whether TREM2 expression is relevant in human AMD, we analyzed flow cytometry data from the same cohort of human donors used for generating the scRNA-seq in this study (**Fig. S6E, Table S2**). We observed that the frequencies of TREM2^+^ myeloid cells (CD45^+^CD11B^+^) were elevated in the RPE/choroid tissues from AMD donors, while the frequencies of CD45^+^ and CD11B^+^ cells remained relatively unchanged (**Fig. 6D-6G, Fig. S6F**). From our scRNA-seq dataset, we can conclude that the identity of TREM2+ myeloid cells are likely microglia. Moroever, we found that the frequencies of TREM2^+^ myeloid cells are strongly correlated with AMD progression (**Fig. 6H**), further supporting our results for subretinal GAL3^+^CD68^+^ histologically. Therefore, our collective findings using orthoganoal approaches implicate the involvement of GAL3-TREM2 in human AMD. Of note, we attempted to immunolabel for TREM2 histologically in retinal sections with anti-human TREM2 antibodies (R&D AF1828 and LSBio LS-B16999), but without success, as others have reported similar technical issues with formalin-fixed and paraffin-embeded CNS tissues (*61, 62*).

## DISCUSSION

Our results uncovered a common population of microglia in the subretinal space that restrict disease progression across multiple distinct mouse models of retinal degeneration. Likewise, we identified microglia with a conserved profile at the transcriptomic, protein, and spatial levels that are present and enriched in the macula of human subjectject with geographic atrophy. As the accumulation of subretinal macrophages has been documented in many, if not most, retinal degenerative diseases (*33–36, 63, 64*), the protective subretinal microglial signature we defined may represent a general response in these disease settings. Although the mouse models used in our study may not represent all clinical aspects of human AMD, our data revealed over 87% DEGs shared between mice and humans, and overlapped functional inferences (such as phagocytosis and lipid activity). Hence, our findings may lay the molecular foundation for using mouse models to understand subretinal microglial responses in human retinal degeneration.

Our results revealed that within the microglial compartment, only subretinal microglia upregulate Gal3 expression, though we do acknowledge that Gal3 can be expressed by non-microglial cells, such as infiltrated monocyte-derived macrophages, astrocytes, and retinal Müller glia in the CNS disease settings (*65–67*). Indeed, several studies using *Lgals3* global KO mice had reported that Gal3 act pathologically in models of neurodegeneration (*44, 68, 69*) and promotes loss of retinal ganglion cells in glaucoma (*70*). The crux of our experiments explicitly focused on microglia via the use of *Cx3cr1^CreER^* mice, which helped identify the isolated roles of Gal3 in microglial-mediated phagocytosis of dead photoreceptors and RPE protection. By contrast, one recent study reported Gal3 inhibition appear neuroprotective and resulted in increased retinal thickness in the LD setting (*71*). However, their increased retinal thickness may be due to deficient phagocytic clearance of dead photoreceptors as identified in our study, and some of the results may be confounded by mouse genetic background. Also, we cannot exclude the possibilities that monocyte-derived cells may be present in the outer retina in AMD (*72*) and that subretinal cells may have neuroinflammatory roles (*37*). Hence, further investigation would be needed to understand Gal3-mediated functions in microglial versus non-microglial cells in different neurodegenerative contexts.

Mechanistically, Gal3-mediated protection required upstream Trem2-signalling. The latter, we showed, regulated microglial migration to the subretinal space and induced Gal3 expression. Trem2-mediated signaling may be particularly important for the subretinal protection. Indeed, we found upregulated Trem2 protein expression and Syk expression in subretinal microglia. Moreover, 4D9 agonistic antibodies dominantly localize with subretinal microglia, likely due to increased Trem2 expression by these subretinal cells. Our data also corroborate the recent findings on neuroprotective roles of microglia-derived Syk in neurodegenerative diseases (*73, 74*).

Trem2, a central lipid sensor by microglia (*49*), could mediate neuroprotection via increased phagocytosis, antioxidant activity and lipid metabolism by subretinal microglia (*6*), and facilitate prompt clearance of dead photoreceptors and cellular debris to support maintenance of RPE homeostasis. Congruently, we showed that conditional genetic depletion of microglia (*6*) or deletion of Gal3 in this study resulted in the accumulation of dead/dying photoreceptor debris in the subretinal space. Separately, as Trem2-Gal3 colocalization was polarized towards the RPE facing aspect of subretinal microglia, subretinal microglia may play a direct role in RPE preservation, but these points require further investigation. Interestingly, the transcriptional signature of these subretinal microglia resembles that of disease associated microglia (DAM) in neurodegeneration (*11, 13*), and also similar with microglial profiles in development (*75, 76*). Consistent with several studies showing that Gal3 is present in this signature (*13, 77, 78*), our study demonstrated that Gal3 is required for Trem2-mediated protection and established their functional association in the retina.

In our study, we also observed elevated levels of soluble Trem2 in retinal degeneration. This elevation in the disease state may result from either increased Trem2 expression by microglia or increased cleavage by secretases (*54, 55*). Indeed, elevated soluble TREM2 was also observed in cerebrospinal fluid from Alzheimer’s patients (*53, 79–82*), and the strategy to bolster TREM2 activity is being tested in a phase II clinical trial. Hence, our findings that subretinal microglia restrict disease progression via Gal3-Trem2 signaling and that this response can be bolstered pharmacologically, may provide a novel focal point for developing potential therapeutic interventions to support photoreceptor and RPE preservation in outer retinal degenerative diseases.

## MATERIALS AND METHODS

### Study Design

The overall goal of this study was to determine the transcriptomics and functional contribution of subretinal microglia in degenerative diseases of the outer retina. We enriched and profiled retinal microglia and other macrophages from four distinct mouse models and identified a common signature of subretinal microglia marked with galectin-3 upregulation. Galectin-3 global knockout mice and microglial-conditional galectin-3 and Trem2 knockout mice were used for loss-of-function assessment of subretinal microglia in multiple degeneration models, while 4D9 anti-Trem2 agonist was used for gain-of-function. Outcomes measured included changes of subretinal Iba1 cells and galectin-3 expression, microglia phagocytosis, changes of soluble Trem2 levels, RPE dysmorphogenesis, photoreceptor death, and vision loss. Analysis of human retinal macrophage include single-cell RNA-seq, immunohistochemistry and flow cytometry. Mice were randomly assigned for experimental groups, and sex was matched among experimental groups. The sample size and replicates of mouse experiments were not predetermined but estimated by literature documentation of similar experiments, while human sample size of human was based on data availability. The sample size and experimental replicates are indicated in the figure legends. Investigators were not blinded for data analysis.

### Mice

All procedures involving animals were approved by the Institutional Animal Care and Use Committee at Duke University, and the procedures were carried out in accordance with the approved guidelines. Wild-type C57BL/6J, *Lgal3^-/-^* (Stock #006338), *Cx3cr1^YFP-CreER^*(Stock #021160), *Trem2^fl/fl^* mice (Stock #029853) were obtained from the Jackson Laboratory. *Rho^P23H^*mice (*38*) were generated as described previously. The frozen sperms of *Lgals3^fl/fl^* mice (*42*) were kindly provided by Bart O. Williams from Van Andel Research Institute and were rederived at Duke University. The rederived mice were further bred into C57BL/6J background. *Cx3cr1^YFP-CreER^*mice were crossed with *Lgals3^fl/fl^* or *Trem2^fl/fl^* to generate the strains for microglia-specific depletion, respectively. Only heterozygous *Cx3cr1^YFP-CreER^*mice were used. If not otherwise stated, mice used included both sex and were at 8-20 weeks of age. All mice herein did not carry *rd8* mutation and were bred and housed at a barrier-free and specific-pathogen-free facility with a 12 h light/12 h dark cycle at Duke University.

### Human autopsy eyes

The metadata of human donors and autopsy eyes were included in **Table S2**. The use of autopsy eyes for research was approved by the Institutional Review Board at Duke University. Due to the lack of ophthalmic clinical history in most cases, the diagnosis of AMD was made postmortem. Following the removal of the superior calotte, postmortem fundus examination and color photography, the eyes were embedded in paraffin and sectioned at 5-μm thickness. Hematoxylin and eosin, periodic-acid Schiff, and immunostained macular sections were evaluated for the presence of AMD and graded using the AMD grading system by Sarks. Eyes with other detectable macular pathology or with glaucoma were excluded.

### Light damage model

Light damage of mice was induced as previously described. Briefly, mice were adapted in darkness overnight, and eyes were dilated with 1% atropine sulfate (Bausch & Lomb) and 10% phenylephrine hydrochloride (Paragon BioTeck). Mice were then placed in a reflective container with a cool white-light LED light source (Fancierstudio), which was placed above the container with 65,000 lux adjusted using an illuminance meter. After 6 h exposure for *Cx3cr1^YFP-CreER^* mice or 8 h for other C57BL/6J mice, the mice were returned to the housing facility with normal lighting and bred for additional five days before experiments.

### Sodium iodate model

Two-month-old mice were administrated a single dose of sodium iodate (Sigma-Aldrich, 15 mg/kg body weight) via intraperitoneal injection. After 5 days, mice were euthanized, and retinas were collected for analysis.

### Conditional depletion in microglia

*Cx3cr1^YFP-CreER/+^*; *Lgals3^fl/fl^* mice*, Cx3cr1^YFP-CreER/+^*; *Trem2^fl/fl^* mice, or *Cx3cr1^YFP-CreER/+^*control mice were intraperitoneally injected with tamoxifen (Sigma-Aldrich, 75 mg/kg) twice with one day in between injections. To target microglia only, after tamoxifen pulse, mice were rested for four weeks before experiments, which spared depletion in monocytes and monocyte-derived cells (*21*).

### Immunohistochemistry

Mice were euthanized by CO_2_ asphyxiation immediately before tissue harvest. Eye tissues were dissected to remove corneas, lens, irises/ciliary bodies, and optic nerves. Tissues were fixed in 4% PFA in PBS for 20 min to 1.5 h at room temperature or on ice. Tissues were either sequentially cryoprotected in 15% and 30% sucrose and then embedded in optimal cutting temperature compound (Tissue-Tek) for cryosections or separated into neuroretinas and RPE/choroids for flat mounts. Flat mounts were blocked and permeabilized with 5% FBS in PBS supplemented with 0.5% Triton-X100 and 0.5% Tween-20, and sequentially incubated with primary antibodies and appropriate secondary antibodies. Phalloidin conjugated with Alexa 594 (Invitrogen #A12381) was included with secondary antibodies to stain F-actin of RPE cells. Primary antibodies used were as follows: rabbit anti-Iba1 (Wako #019-19741), goat anti-Gal3 (R&D #AF1197), rat anti-Gal3 (Biolegend #125401), sheep anti-Trem2 (R&D #AF1729), mouse anti-rhodopsin (Abcam #ab5417) and rabbit anti-Syk (Abcam #ab40781). Images were acquired using a Nikon A1R confocal laser scanning microscope. A resonant scanner and motorized stage were used to acquire *z*-stacks. Unless otherwise indicated, maximum projections of image stacks were shown.

Human autopsy eyes were fixed in 3.7% neutral-buffered formaldehyde. The detection of Galetin-3 and CD68 was performed at Duke Pathology Core facility using The VENTANA DISCOVERY Ultra automated immunohistochemistry staining system (Ventana Medical Systems). Sections were incubated with primary antibodies: rat anti-human Gal3 (Invitrogen #14-5301-82, 1:50 dilution) and mouse anti-human CD68 (Dako #M0814, 1:400 dilution), following by the secondary antibody incubation and chromogenic detection with DISCOVERY Purple and Yellow kits (Ventana-Roche Diagnostics, #760-229 and # 760-239). These chromogenic dyes are covalently deposited and have unique spectra (*83*) that allow spatial mapping and detection of colocalization using multispectral imaging (*84*). Cells coexpressing Discovery Yellow and Purple appear orange-red (*85*), and coexpression was confirmed using a Nuance 3.0.2 Multispectral Imaging System (PerkinElmer).

### Histology in mice

Euthanized mice were fixed via transcardial perfusion with 2% paraformaldehyde and 2% glutaraldehyde in 0.1% cacodylate buffer (pH = 7.2). The eye tissues were post-fixed in the same fixative for 24 h and processed in a solution of 2% osmium tetroxide in 0.1% cacodylate buffer, following by processing with gradient ethanol from 50% to 100%, propylene oxide, and propylene oxide: epoxy 812 compound (1:1 ratio) under the vacuum. Samples were further embedded in fresh epoxy 812 compound resins at 65°C overnight. Semi-thin cross sections (0.5 μm) across the block were stained with 1% methylene blue.

### Morphological analysis of microglia

The covered area and process length of microglia were quantified as previously described (*86*). Briefly, images of neuroretina or RPE/choroid flat mounts stained with Iba1 were optimized and transformed into binary ones. The covered area of individual microglia was measured using Analyze Particle in ImageJ, and the average covered area per image was calculated as the total areas divided by the number of microglia. To quantify the process length, images were skeletonized and analyzed using the Analyze Skeleton (2D/3D) Plugin in ImageJ. The branch length of individual microglia was summed and divided by the number of microglia. For each mouse model, four mice per group and three images (628.22 μm x 628.22 μm) per mouse n=4 mice were analyzed.

### Quantifications of RPE dysmorphology and subretinal microglia

RPE dysmorphology was quantified as described previously (*6*). RPE flatmounts stained with phalloidin and Iba1 were imaged, and multiplane z-series images were acquired using 20x objective (628.22 μm x 628.22 μm per image). To avoid the confounding effects from the areas around optic nerve head and peripheral iris/ciliary bodies, one random image was acquired in the middle of each RPE quadrant, with a total of four images per RPE flatmouts. RPE cells that exhibited either altered lateral or lost apical F-actin morphology were considered as dysmorphic. Four complete fields per RPE flatmount were assessed. The numbers of abnormal and total RPE cells were counted in each field, and the mean percentage of RPE dysmorphology in four fields was calculated for each mouse. The frequencies of subretinal microglia per field were determined as total cell counts divided by total areas, and the mean frequency of subretinal microglia in four fields was calculated for each mouse.

### TUNEL assay

This assay was performed using *in situ* cell death detection kit (Roche) according to manufacturer’s instruction. Briefly, retinal cross sections were blocked and stained for TUNEL and DAPI. At least three images of each animal were acquired using Nikon A1R confocal microscopy and analyzed using Image J. The frequencies of TUNEL positive cells per 1 mm^2^ in the ONL were calculated.

### Quantifications of phagocytosis by subretinal microglia

Retinal cross-sections were stained with anti-rhodopsin and anti-Iba1 antibodies. Nuclei were counterstained with DAPI. Single planes of confocal scans were used to quantify rhodopsin-positive microglia in the subretinal space. Three images of each mouse were acquired, and the mean percentage of rhodopsin^+^ cells was calculated for each mouse.

### OCT and Fundus Imaging

Mouse eyes were topically dilated with 1% tropicamide and 10% phenylephrine sulphate and anesthetized via intraperitonially injection with a mixture of ketamine/xylazine. The corneas were kept moist with GenTeal^®^ lubricant eye gel (Alcon). Eyes were imaged using Micron IV retinal imaging system (Phoenix Research Labs).

### Electroretinogram (ERG)

ERG was measured as previously described (*6*). Briefly, pupils of dark-adapted mice were dilated with 0.5% tropicamide and 1.25% phenylephrine. and mice were then anesthetized with a mixture of ketamine/xylazine. Scotopic and photopic responses were recorded using an Espion E2 system (Diagnosys) with increasing flash intensities (scotopic: from 2.5 × 10^−5^ to 500 cd·s/m^2^; photopic: from 5 to 500 cd·s/m^2^ with a background light of 25.5 cd·s/m^2^ intensity). Recordings of single flash presentations were measured 1–15 times to verify the response reliability and improve the signal-to-noise ratio, if required.

### Treatment of anti-Trem2 antibodies

*Cx3cr1^YFP-CreER/+^*mice were injected via tail vein with Fc-mutated mAb178 anti-Trem2 (50 mg/kg) or vehicle control for loss-function, or 4D9 anti-Trem2 (50 mg/kg) or isotype for gain-function, right before dark adaptation of light damage.

### Quantifications of human IgG containing microglia

To determine the location of 4D9 antibodies, retinal cross-sections, retinal and RPE/choroid flat mounts were stained with DyLight^TM^ 594 donkey anti-human IgG and anti-Iba1 antibodies were imaged with Nikon A1R confocal microscopy. The retinas from mice subjected to LD without 4D9 treatment were used as negative controls for autofluorescence. The human IgG^+^ microglia were counted on the RPE and in the inner retina, and the percentages were shown.

### ELISA of vitreous and retinal fluids

Vitreous fluids were collected from euthanized mice using a Hamilton syringe with a 30-gauge needle and immediately mixed with proteinase inhibitors. For retinal fluids, retinas were dissected in a dry dish and then incubated on ice for 10 min with 50 μl PBS per retina supplemented with proteinase inhibitors. After centrifuging at 14,800 g for 5 min, fluid samples were collected. ELISA of soluble Trem2 was measured using DuoSet^®^ Ancillary Reagent Kit 2 (R&D) according to the manufacturer’s instructions. Recombinant Mouse Trem2 (R&D #1729-T2) was used to generate a standard curve. Capture antibody of anti-mouse Trem2 (R&D #AF1729) and detection antibody of biotinylated anti-Trem2 (R&D #BAF1729) were used at 0.4 μg/ml and 0.1 μg/ml, respectively. Biotinylated antibodies were detected using streptavidin-HRP (Biolegend #405210).

### Single cell RNA-sequencing

Mouse retinas were dissected from 5 males of each model, including 2-month-old naïve wild-type mice, 2-month-old mice of sodium iodate model, 2-month-old *Rho^P23H/+^* mice, and 2-year-old wild-type mice. Retinas of each model were pooled and digested in 1.5 mg/ml collagenase A and 0.4 mg/ml DNase I (Roche) for 45 min at 37°C with agitation. Single-cell suspensions were generated by passing through 70 μm filters and sequentially stained with APC anti-mouse CD45 (Biolegend #103111) and propidium iodide (Sigma) for viability. Viable CD45^+^ single cells were collected by Fluorescence Activated Cell Sorting (FACS). 10x Genomics Single Cell 3′ chemistry (v2) was used to generate Gel Bead-In Emulsions (GEM), and perform post GEM-RT cleanup, cDNA amplification, as well as library construction.

Eye tissues from human donors were recovered within 8 hours of death and then dissected to separate neuroretinas and RPE/choroids. Neuroretinas were then homogenized using douncers and RPE/choroids were digested with collagenases A and DNase I for 1 hour at 37°C with agitation, respectively. Single-cell suspensions were generated by passing through 70 μm filters, processed with debris removal solution (Miltenyi Biotec) and then frozen and stored in Recovery Freezing Medium (Thermo Fisher). Frozen cells were thawed in 5% Fetal and sequentially stained with viability dye eFluor 450 (eBioscience #65-0863-14), BV785 anti-human CD45 (Biolegend #304048), BV510 anti-human CD11B (Biolegend #562950) and processed with single-cell multiplexing kit (BD). Samples were also stained with APC anti-human TREM2 (R&D #FAB17291A) for the downstream analysis. CD45^+^CD11B^+^ cells were sorted and loaded into BD Rhapsody single-cell analysis system. The cDNA libraries were prepared using BD Rhapsody whole transcriptome analysis amplification kit.

Agilent DNA 4200 Tapestation assay was used for quality control. Libraries were pooled and sequenced to target 50,000 unique reads per cell using an Illumina NextSeq (high run type) for mice and an Illumina NovaSeq6000 (S1 flow cell) for human and with the read length of 75 base-pairs and paired-end.

### Analysis of *de novo* scRNA-seq data

Mouse and human raw sequencing data were initially processed with Cell Ranger pipelines for 10x Genomics and Seven Bridges pipelines for BD Rhapsody, respectively. Briefly, FASTQ files were generated by demultiplexing and further aligned to the mouse genome reference mm10 and the human genome reference GRCh38, respectively. Feature barcode processing and unique molecular identifier (UMI) counting were then performed according to the standard workflow. The following criteria were applied as quality control using Seurat (*87*) (v4): cells that had fewer than 200 UMI counts or genes that were expressed by fewer than 3 cells were removed from further analysis. Mouse cells that had more than 5,000 UMI counts or greater than 20% of mitochondrial genes were also excluded, while doublets of human cells were identified and removed after clustering analysis.

Data integration was performed using Seurat (*87*). Specifically, the mouse datasets were integrated with a previously published retinal microglial dataset from light damage model (*6*), and the datasets of human neuroretina and RPE/choroids were also integrated before clustering analysis. After filtering, top 2,500 and 2,000 features were selected to identify the anchors for mouse and human datasets, respectively. Top 30 PCs were used to generate UMAP clustering. To identify the conserved marker genes of subretinal microglia in mice, differential gene analysis was performed in each model, and then the overlapped markers were selected.

Gene module scores of homeostatic microglia, subretinal microglia, perivascular macrophages and monocyte-derived macrophages were calculated as previously described (*57*). For each population, top 200 differentially expressed genes ranked by fold change and identified in a mouse model were used generate the module score with AddModuleScore function in Seurat. Data were visualized with DotPlot.

Pathway enrichment analysis were performed using top 200 shared upregulated genes that were ranked by average fold change of subretinal microglial clusters from mice and human donors. Gene ontology database with biological process was used (http://geneontology.org). A pathway was considered significantly over-presented with FDR < 0.05.

### Integration and analysis of independent AMD scRNA-seq datasets

The scRNA-seq datasets from another two independent AMD studies were downloaded with the accession number GSE183320 (*88*) and GSE203499, respectively. Clustering analysis was first performed to extract myeloid cells by *AIF1* expression in these two datasets. The myeloid cells were further integrated with our *de novo* dataset using Seurat. Top 3,000 features were used to identify the anchors for integration and top 30 PCs were used to generate UMAP clustering.

The label transfer was performed with the default setting of Seurat tutorial. Our *de novo* dataset was used as the reference, and the other two datasets were used as queries. The *RHO^high^* cluster unique present in one AMD donor were excluded from the label transfer. Based on the metadata availability, the frequencies of subretinal microglia cluster relative to all myeloid cells in the whole or the macular RPE/choroid were calculated for each donor.

### Flow cytometry analysis

Data of human RPE/choroid tissues were collected using BD FACSAria III Cell Sorter and analyzed using FlowJo software (version 10.7.2). Control human blood was used for gating viability, all immune cells, and myeloid cells. CD45^+^CD11B^-^ cells from human blood were used as a negative control for gating TREM2^+^ cells in RPE/choroid tissues.

### Statistical analysis

Data are presented as means ± standard errors. Normal distribution and homogeneity of variance were tested before applying any parametric analysis, and data transformation was performed when needed. If the assumptions of a parametric test cannot fit, a non-parametric test was used. Depending on the research questions, one-tailed or two-tailed tests were used. A student’s t-test or Mann-Whitney test was used for two group comparisons. For multiple comparisons, one-way or two-way ANOVAs followed by Tukey’s post-hoc test were used. For correlation analysis, Spearman’s correlation coefficient was used for rank-ordered data. All non-sequencing experiments were repeated at least twice. A p-value less than 0.05 is considered statistically significant. All statistical data were analyzed using GraphPad Prism.

## Supplementary Materials

This PDF file includes:

Figs. S1 to S6.

Tables S1 to S3.

## Acknowledgments

We thank Kathryn Monroe at Denali Therapeutics for providing the modified 4D9 antibodies. We thank Bart O. Williams for providing frozen sperms of *Lgals3^fl/fl^* mice. We acknowledge technical assistance from Duke core facilities.

## Author contributions

Conceptualization: CY, DRS; Data acquisition: CY, EML, RM, JK, SL, YC, KS, ADP; Data analysis: CY, EML, SL, ADP, DRS; Data interpretation: CY, EML, YC, KS, CBR, ADP, MC, CH, DRS; Visualization: CY, DRS; Writing – original draft: CY, DRS; Writing – review & editing: all authors; Supervision: EML, ADP, MC, CH, DRS; Project administration: DRS.

## Data and materials availability

Two human and mouse scRNA-seq datasets generated by this study have been deposited in the Gene Expression Omnibus (GEO) with the accession numbers GSE208434 and GSE195891, respectively. The retinal microglial dataset from light damage model was under the accession number GSE126783. The other two datasets of human AMD were downloaded with the accession number GSE183320 and GSE203499, respectively. All the analytic scripts are available upon request. All other data needed to evaluate the conclusions in this paper are available in the paper or the Supplementary Materials.

---

**Fig. S1.**
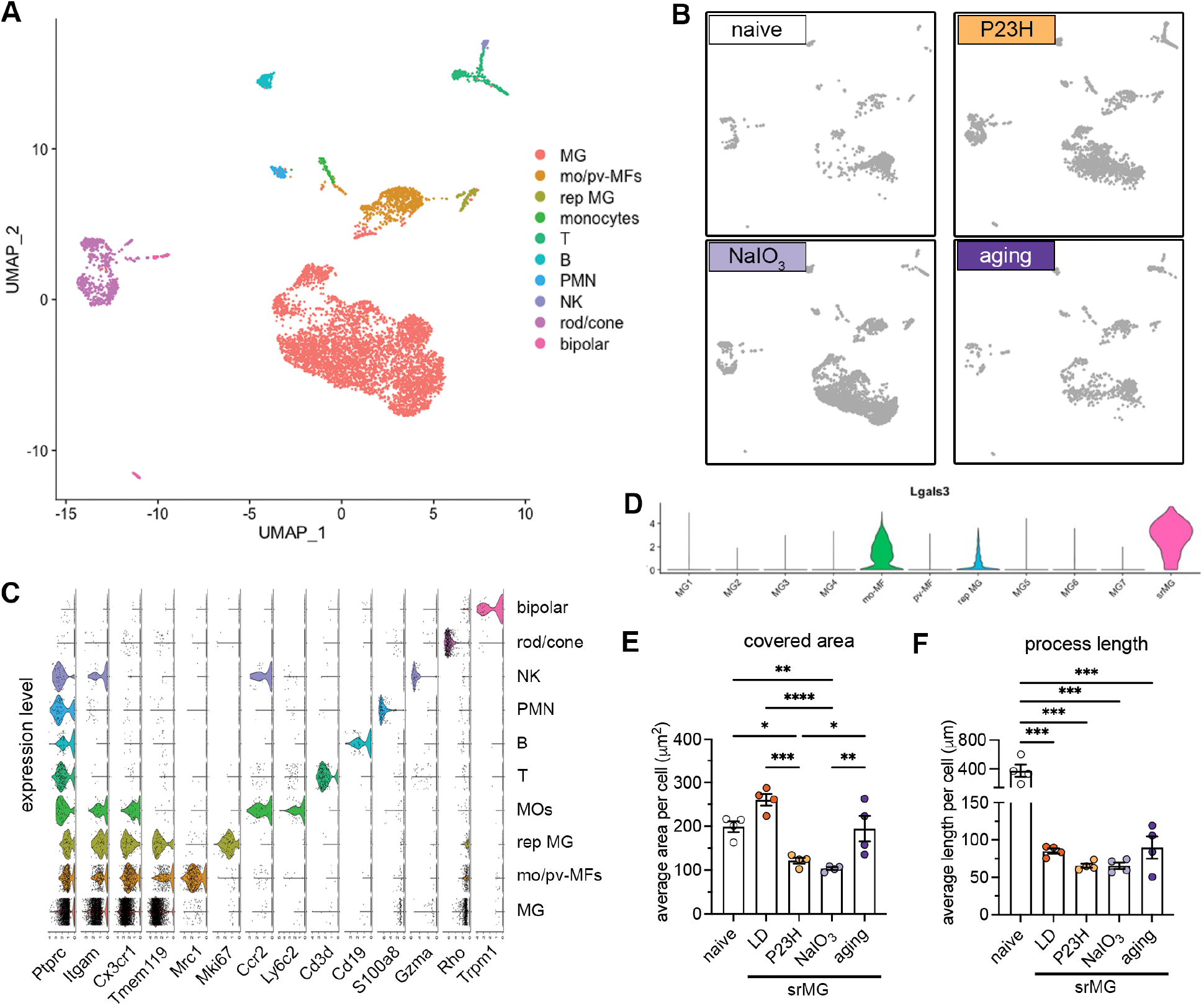
scRNA-seq and morphological analysis of subretinal microglia across mouse models of outer retinal degeneration. (**A** and **B**), UMAP plots showing retinal CD45^+^ cells collected from naïve mice, NaIO_3_ mediated RPE injury model, P23H model, and advanced aging model as indicated. (**C**) Violin plots showing marker expression for each cluster. mo-MF, monocyte-derived macrophages; pv-MF, perivascular macrophages; rep MG, replicating microglia. (**D**) Violin plots showing *Lgals3* expression across all macrophage clusters. srMG, subretinal microglia. (**E** and **F**) Quantifications of covered area and process length in naïve microglia from the inner retina and subretinal microglia from four mouse models of retinal degenerations (n=4 mice per group). *: p<0.05; **: p<0.01; ***: p<0.001 (one-way ANOVA with Tukey’s post hoc test)

**Fig. S2.**
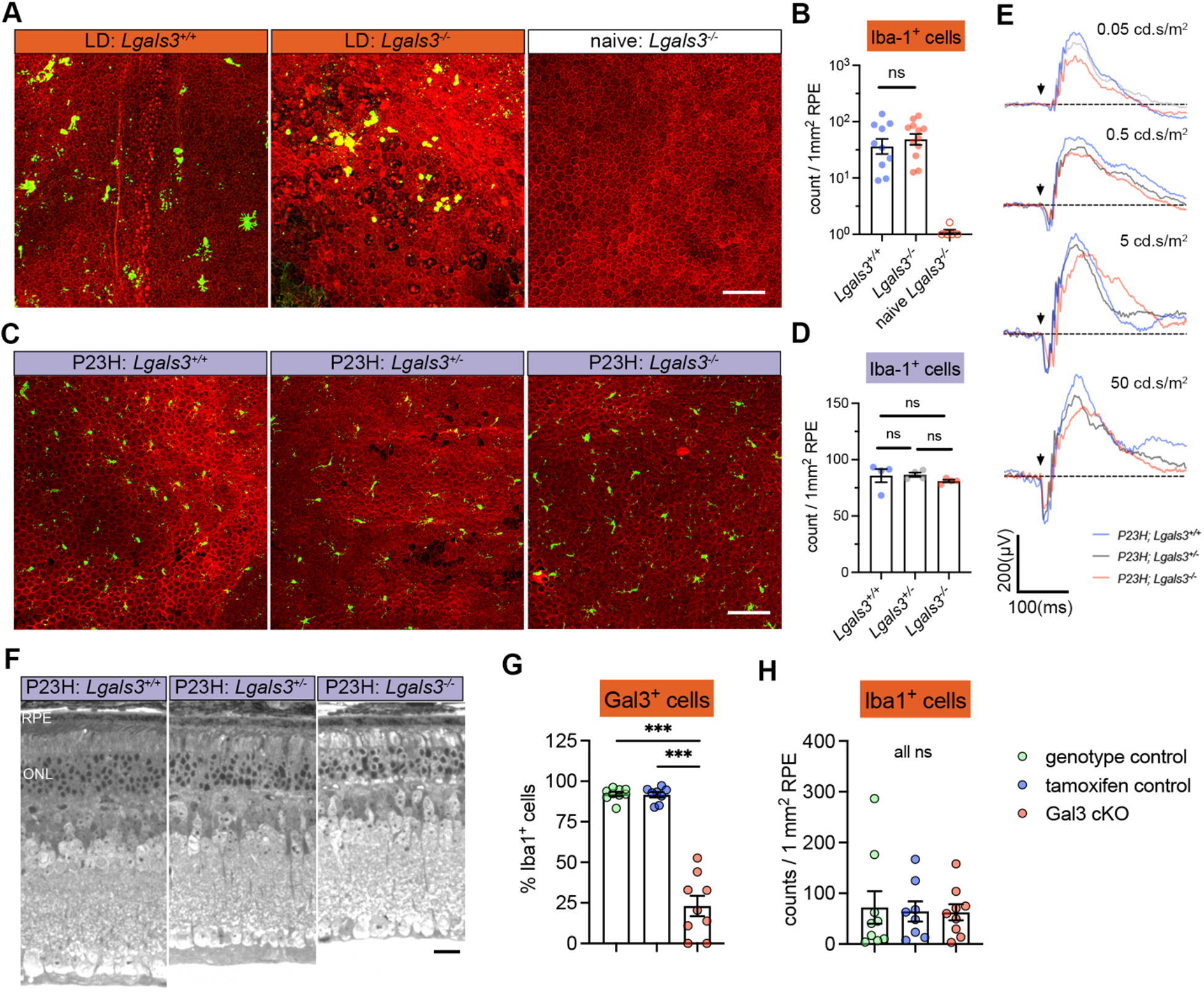
Contributions of Gal3 to disease-related retinal pathology and Iba1^+^ cell abundance in the subretinal space. (**A**) Iba1 (green) and phalloidin (red) staining in RPE flatmounts from LD-subjected mice as indicated. (**B**) Quantifications of subretinal Iba1^+^ cells as shown in A. (**C**) Iba1 (green) and phalloidin (red) staining in RPE flatmounts from P23H mice as indicated. (**D**) Quantifications of subretinal Iba1^+^ cells as shown in C. (**E**) Examples of ERG responses at different flash intensities as indicated. (**F**) Representative retinal cross sections of WT, *Lgal3^+/-^*and *Lgal3^-/-^* in P23H mice. (**G** and **H**) Quantifications of Gal3 depletion efficiency (G) and frequencies of subretinal Iba1^+^ cells (H) in Gal3 cKO mice (n=9) compared with genotype control mice (n=9) and tamoxifen control (n=8). Scale bars: 100 μm. Data were collected from 2-3 independent experiments. ***: p<0.001; ns: not significant (one-way ANOVA with Tukey’s post hoc test).

**Fig. S3.**
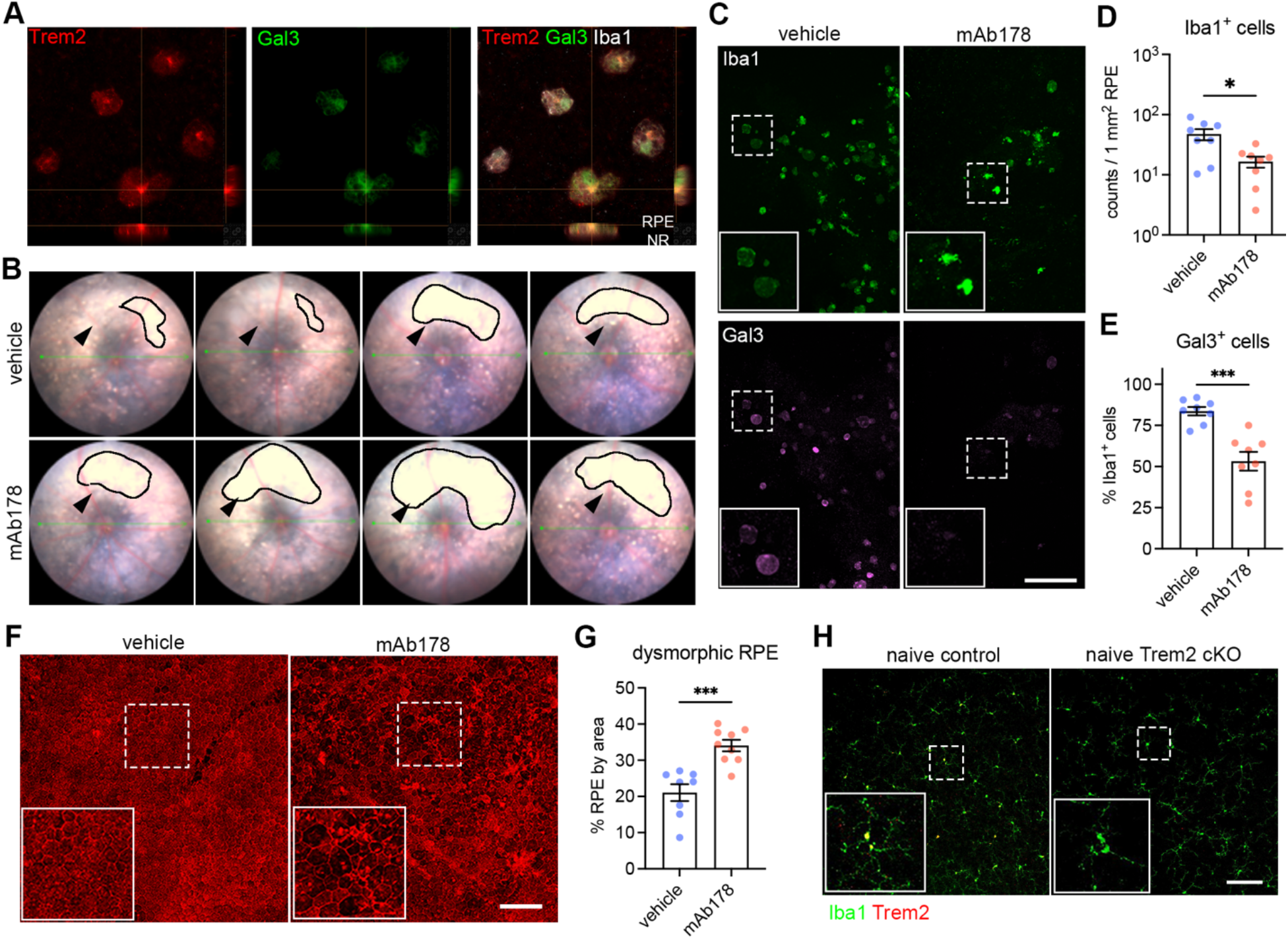
Regulation by Trem2 signaling in subretinal microglia. (**A**) Split views of confocal scans showing the colocalization of Trem2 (red) and Gal3 (green) in the subretinal microglia. Lines indicate the RPE-facing and neuroretina (NR)-facing aspects as indicated. (**B**) Fundus images showing increased subretinal white lesions in anti-Trem2 mAb178 treated mice in LD as indicated by arrows. Images of 4 individual mice per group are shown. (**C**) Images of Iba1 (green) and Gal3 (magenta) staining in subretinal microglia between control and mAb178-treated mice in LD. Scale bar: 100 μm. (**D** and **E**) Quantifications of Iba1^+^ cells and Gal3^+^ cells between control and mAb178 (n=8 per group). (**F**) Images of phalloidin staining in RPE flatmounts from control and mAb178 treated mice in LD. Scale bar: 100μm. (**G**) Quantifications of dysmorphic RPE cells between control (n=8) and mAb178 (n=9) treated mice. (**H**) Images of Iba1 (green) and Trem2 (red) in microglia from the inner retina of naïve control and Trem2 cKO mice. Scale bar: 50μm.

**Fig. S4.**
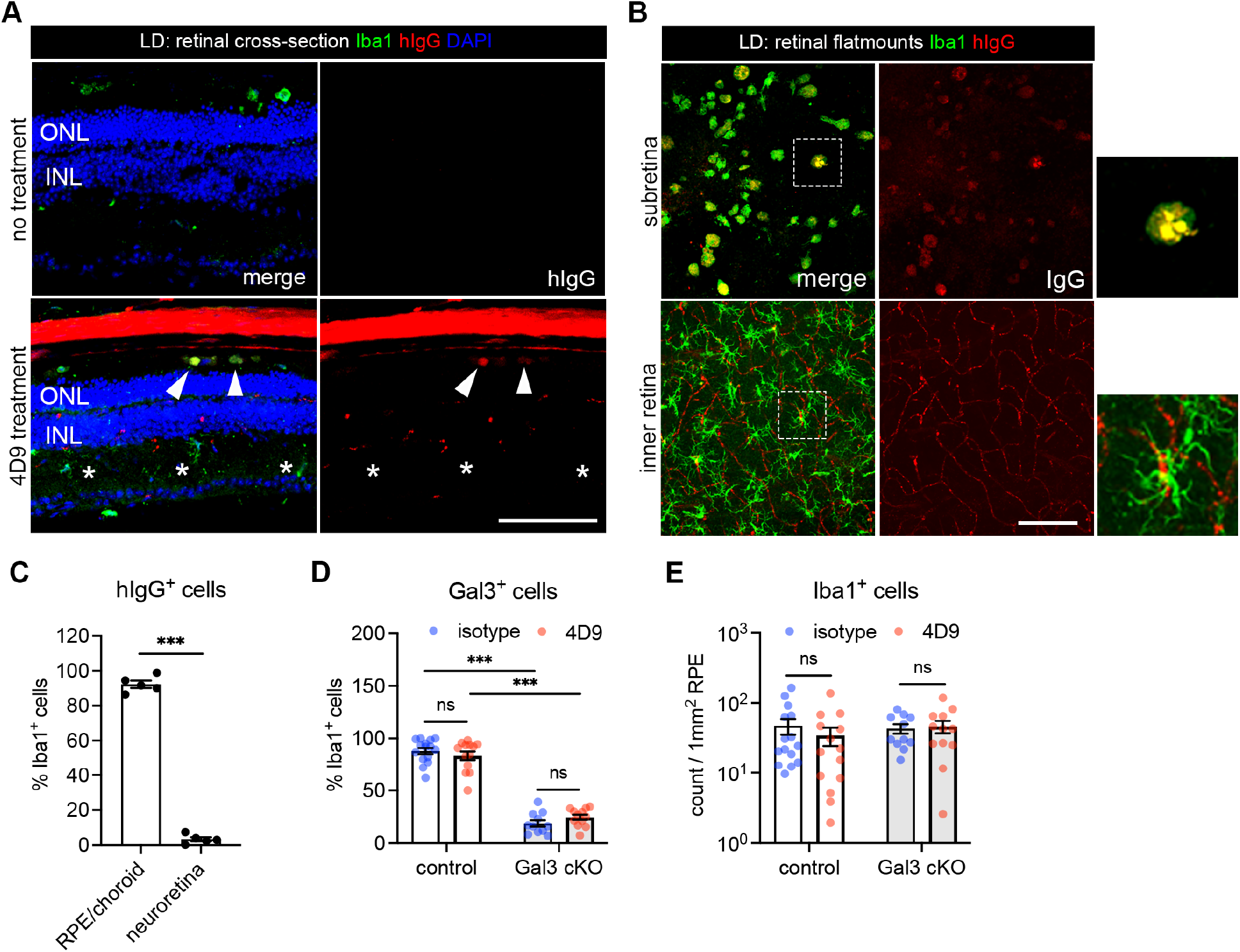
Subretinal microglia with 4D9 treatment. (**A**) Staining of human IgG (red) and Iba1 (green) in retinal cross sections collected from mice with or without 4D9 treatment in LD. The hIgG is used to trace 4D9 antibodies, which outlines retinal vasculatures in 4D9 treated mice. Arrows indicate the presence of 4D9 antibodies in the subretinal microglia, while asters indicate the absence of 4D9 antibodies in microglia from the inner retina. (**B**) Human IgG (red) and Iba1 (green) staining in RPE and neuroretina flatmounts collected from mice treated with 4D9 antibodies in LD. (**C**) Quantifications of hIgG^+^ microglia in the subretinal space and neuroretina. (**D** and **E**) Quantifications of Iba1^+^ cells and Gal3^+^ cells between control and Gal3 cKO mice treated with either isotype or 4D9 (n=13 per group). Scale bars: 100 μm. Data were collected from 2-4 independent experiments. ***: p<0.001; ns: not significant (unpaired Student’s t-test: C; two-way ANOVA with Tukey’s post hoc test: D and E).

**Fig. S5.**
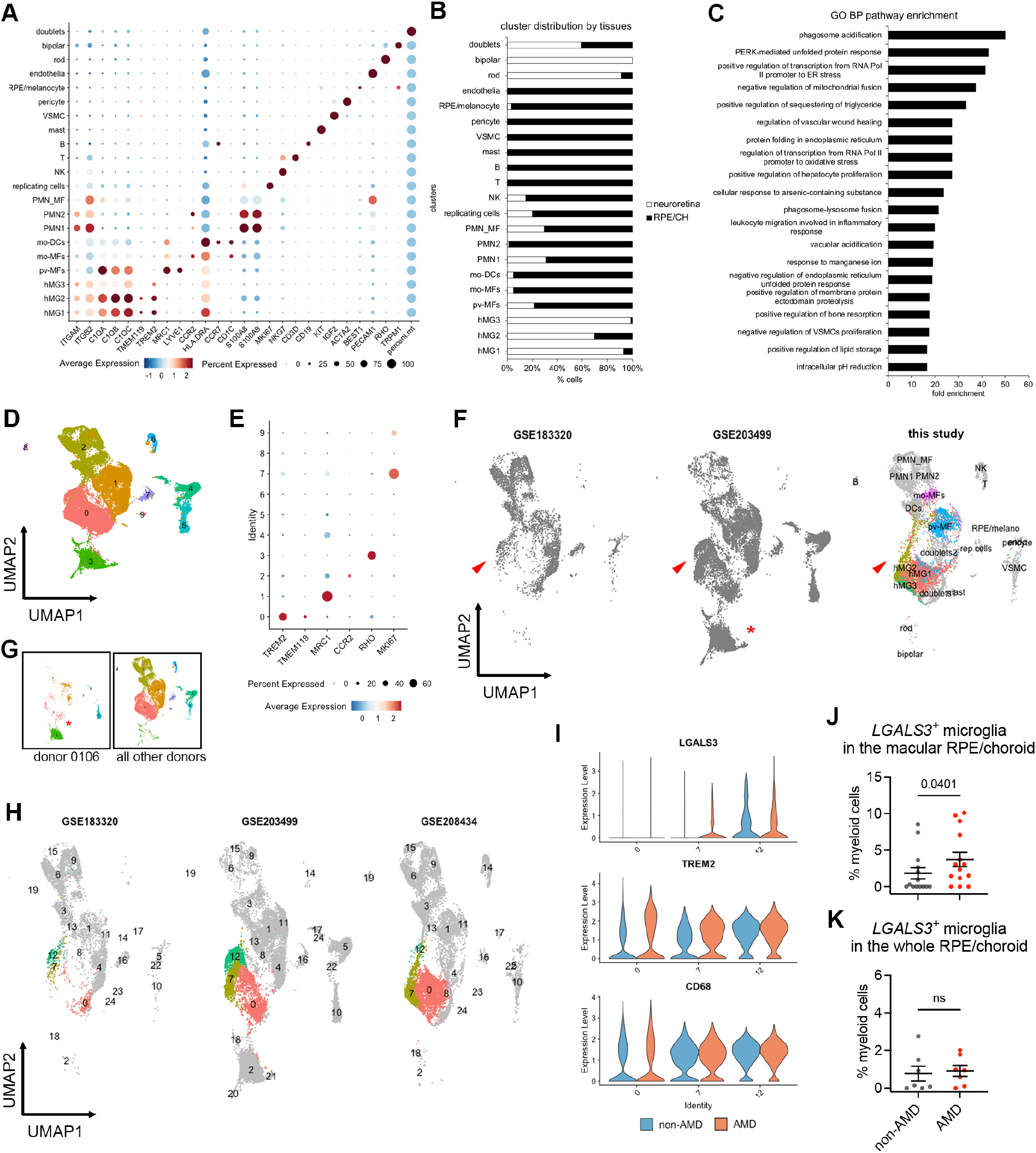
ScRNA-seq analysis of myeloid cells from human non-AMD and AMD donors. (**A**) Marker expression of all human clusters. hMG, human microglia; mo-MFs, monocyte-derived macrophages; pv-MFs: perivascular macrophages; mo-DCs, monocyte-derived dendritic cells; VSMC, vascular smooth muscle cells. (**B**) Distribution of clusters by neuroretina and RPE/choroid tissues. Cell number of clusters was normalized to the total counts per tissue. (**C**) Pathway enrichment analysis of subretinal microglia with top 200 shared up-regulated genes. Top significant pathways sorted by false discovery and ranked by fold enrichment are shown. (**D**) UMAP plot showing integrated clustering analysis of three independent human AMD datasets. Data are shown with low resolution to reveal major cell types. (**E**) Dot plot showing the marker expression of major macrophage clusters. Cluster 3 is enriched with *RHO* expression. (**F**) UMAP plots showing the presence of hMG2 cluster in all three scRNA-seq datasets as indicated by arrows. (**G**) UMAP plots showing the enrichment of cluster 3 in donor 0106_nAMD. (**H**) UMAP plots showing clustering analysis with high resolution by each dataset and comparable heterogeneity of microglia (cluster 0, 7 and 12). As dataset GSE183320 does not contain neurosensory retina tissues, few cells of major homeostatic microglia (cluster 0) are observed in this dataset. (**I**) Violin plots showing the expression of *LGALS3*, *TREM2* and *CD68* by microglial clusters between non-AMD and AMD donors. Both cluster 7 and 12 show *LGALS3* upregulation as hMG2 cluster identified in this study. (**J** and **K**) Quantifications of LGALS3^+^ microglial clusters (7 and 12) in the macular and whole RPE/choroid tissues between non-AMD and AMD donors. Data were from three independent datasets and compared using Mann-Whitney test. P-values are shown. ns: not significant.

**Fig. S6.**
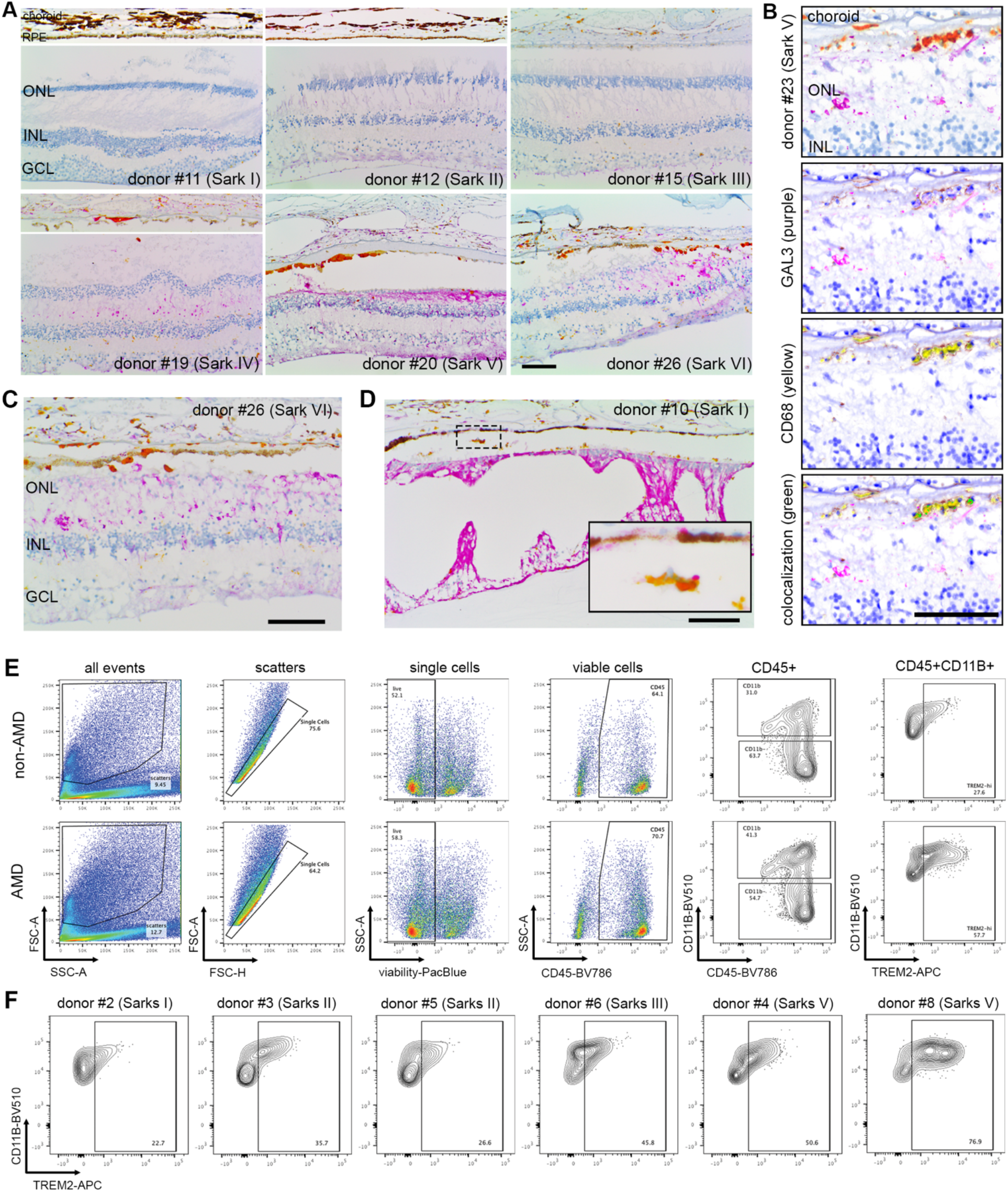
Validation of GAL3 and TREM2 expression by subretinal myeloid cells in human AMD. (**A**) Images of GAL3 (purple) and CD68 (yellow) co-staining in the macula region of retinal sections from human donors categorized by Sark grades (I-VI). The macular neurosensory retinas of some subject eyes exhibited fixation-related artifactual detachment. In these subjects, separate images of RPE/choroid tissues are shown. Scale bar: 100μm. ONL and INL, outer and inner nuclear layers. GCL, ganglion cell layer. (**B**) Spectral imaging of GAL3 and CD68 co-staining in the geographic atrophy from donor #23 with advanced AMD (Sarks V). Unmixed purple spectrum (GAL3) and yellow spectrum (CD68) are shown. The areas of colocalized spectra are highlighted in green. Scale bar: 50μm. (**C** and **D**) Images showing the presence of subretinal GAL3 (purple) and CD68 (yellow) double positive cells in the areas with photoreceptor loss and preserved RPE in the transitional area of the macula from an AMD donor (C) and in the age-related peripheral degeneration of a non-AMD donor (D). Scale bars: 100μm. (**E**) Gating strategy of flow cytometry analysis. CD45^+^CD11B^+^ cells and CD45^+^CD11B^-^ cells from control blood were used to determine the gating of TREM2^+^ cells. Concatenated plots are shown for non-AMD and AMD. (**F**) Flow contour plots of individual donors showing increased percentage of TREM2^+^ myeloid cells in AMD.

